# scGenePT: Is language all you need for modeling single-cell perturbations?

**DOI:** 10.1101/2024.10.23.619972

**Authors:** Ana-Maria Istrate, Donghui Li, Theofanis Karaletsos

## Abstract

Modeling single-cell perturbations is a crucial task in the field of single-cell biology. Predicting the effect of up or down gene regulation or drug treatment on the gene expression profile of a cell can open avenues in understanding biological mechanisms and potentially treating disease. Most foundation models for single-cell biology learn from scRNA-seq counts, using experimental data as a modality to generate gene representations. Similarly, the scientific literature holds a plethora of information that can be used in generating gene representations using a different modality - language - as the basis. In this work, we study the effect of using both language and experimental data in modeling genes for perturbation prediction. We show that textual representations of genes provide additive and complementary value to gene representations learned from experimental data alone in predicting perturbation outcomes for single-cell data. We find that textual representations alone are not as powerful as biologically learned gene representations, but can serve as useful prior information. We show that different types of scientific knowledge represented as language induce different types of prior knowledge. For example, in the datasets we study, subcellular location helps the most for predicting the effect of single-gene perturbations, and protein information helps the most for modeling perturbation effects of interactions of combinations of genes. We validate our findings by extending the popular scGPT model, a foundation model trained on scRNA-seq counts, to incorporate language embeddings at the gene level. We start with NCBI gene card and UniProt protein summaries from the genePT approach and add gene function annotations from the Gene Ontology (GO). We name our model “scGenePT”, representing the combination of ideas from these two models. Our work sheds light on the value of integrating multiple sources of knowledge in modeling single-cell data, highlighting the effect of language in enhancing biological representations learned from experimental data.

## 1 Introduction

Foundation models have seen tremendous success due to their ability to learn meaningful representations from large amounts of data. Inspired by natural language processing [1–6] and computer vision [7], foundation models have become a cornerstone modeling choice for many biological areas as well [8–11]. Single-cell biology in particular is an area where these models have started to be widely applied, due to the availability of large single-cell RNA sequencing datasets and repositories that make the data easily accessible [12]. A number of foundation models for single-cell biology have been developed showing great promise, including scGPT[8], GeneFormer[11], UCE[10], scMulan[13].

Generally, these models learn gene and gene counts representations by modeling the biological variation in scRNA-seq experimental data, using expression counts as a modality for gene and ultimately, cell representations. At the same time, there are other modalities that can be considered when modeling genes. Language in particular is a representation that foundation models like genePT [9] have been exploring. In this paradigm, gene representations are aggregations of various snippets from the scientific literature. This approach is well-justified by the fact that historically findings from experimental data have been disseminated by and to the scientific community through published research articles. This means that there is a large body of knowledge on biological data locked in the scientific literature. This information can be helpful when modeling, guiding or augmenting representations learned from biological entities, such as genes. Moreover, this information can be aggregated in meaningful ways through large language models, which can place textual information within the greater human knowledge context through embeddings.

In this paper, we explore the effect of representing genes through these two modalities when modeling single-cell data: biological representation learned from experimental scRNA-seq counts, and language representation, abstracted as gene information from different sources from the scientific literature. In particular, we are interested in the effect of using these two modalities to model genetic perturbations - the effect of perturbation on transcriptional phenotypes (up- or down-regulation of genes in response to the perturbation). The task of perturbation prediction is usually tackled from two angles: either foundation models that learn representations from data at scale during pretraining, and are then fine-tuned on perturbation prediction as a downstream task (e.g. scGPT[8]), or specialized models that learn from smaller scale, more curated data and embed specific task-related structured information directly into the model architecture (e.g. GEARS [14]). The research questions that guide our efforts are the following:

1. Can we build models powerful enough to learn the structured biological information specific to specialized tasks without having to hardcode it into the model architecture?
2. Will a multimodal approach using language to complement experimental data help us get there?
3. Will curating the knowledge we put into the model have a significant effect?

To explore our hypotheses, we take a popular pre-trained foundation model trained exclusively on scRNA-seq counts, scGPT [8], and inject language into the model architecture at the gene level. Each gene gets a textual representation through LLM embeddings aggregating gene information from various knowledge sources. Inspired by genePT [8], we start by experimenting with NCBI gene card summaries [15] and NCBI gene card summaries combined with UniProt [16] protein summaries. We build on top of this work by testing other sources of information from the scientific literature such as molecular function, biological process and cellular component gene annotations from the Gene Ontology Knowledgebase [17, 18], which we embed using GPT-3.5 [19].

In our analyses, we find that:

1. Textual gene representations provide additive and complementary value to biologically learned gene representations in modeling single-cell perturbations.
2. Textual gene representations are not as powerful as biologically learned gene representations, but provide useful information.
3. Different types of scientific knowledge provide different types of prior information. In the datasets we tested GO Ontology Annotations capturing subcellular localization help the most in single-gene perturbations, and NCBI protein summaries provide the highest value for modeling perturbation effects of interactions of combinations of genes.
4. By carefully curating the auxiliary language-encoded data we introduce into the scGPT transcriptomic foundation model, we can reach and sometime surpass the performance of bespoke models that hard-code structured information explicitly into the model architecture.

We leverage the scGPT and genePT foundation models and show the additive and complementary effect of the two modalities they use (scRNAseq counts, and language). We call our collection of models scGenePT, a testament to the two models we are inspired by and build on top of. Our work casts light on the tremendous potential of multi-modal models, highlighting the value of language in particular to enhance representations learned from experimental data.

## 2 Methods

### 2.1 Perturbation Modeling

There are many types of perturbation - genetic perturbation (e.g. CRISPR), chemical perturbation (e.g. drug treatment), environmental perturbation, infections (e.g. viruses), natural allelic variation (e.g. genetic mutation). For the purpose of our work, we are focusing on modeling genetic perturbations - the effect of perturbing specific genes on the gene expression profile of a cell. Essentially, we are measuring the transcriptome after perturbation. Assume we have a list of *N* genes *G*_*all*_ = [*g*^1^, *g*^2^, …*g*^*N*^] and *M* cell observations *C* = [*c*^1^, *c*^2^, …*c*^*M*^]. For each cell observation *c*^*i*^, we have the corresponding gene expression sequence over the *N* genes:

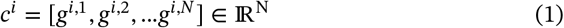

We would like to predict the effect of perturbing specific genes in the initial set *G* on the cell profile. Empirically, we choose a set of *K* genes *G*_*perturb,K*_ = {*g*^1,∗^, *g*^2,∗^, …, *g*^*K*,∗^} ∈ *G*_*all*_ to perturb and for each cell observation *c*^*i*^, we would like to predict the gene expression values post-perturbation:

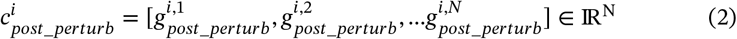

In this work, we explore the following types of perturbations:

- **K = 1**: single-gene perturbations: one specific gene is perturbed
- **K = 2**: two-gene perturbations: two genes are perturbed

### 2.2 Model Architecture

We test our hypotheses by modifying the scGPT architecture to inject language into the model. In scGPT, gene representations include gene expression counts, gene tokens and condition tokens (which in the case of perturbation prediction are perturbation tokens). For scGenePT, each gene gets an additional representation, a gene language representation. All of these different gene representations get added to obtain the final gene representation. An example of how this looks like for gene FOSB can be seen in Figure 3 - the gene gets representations learned from experimental data, and extra information through the language representation (which in the illustrated case is the NCBI Gene Card Summary of the gene).

**FIGURE 1.**
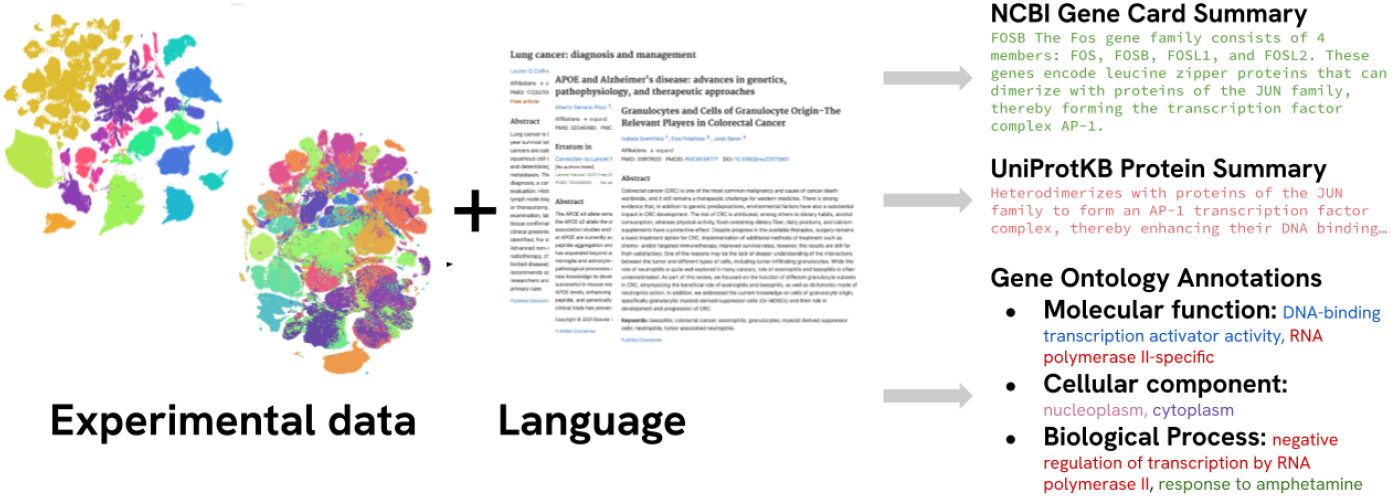
Genes can have representations learned from different modalities: **experimental data**, (e.g.scRNAseq counts) or **language** - through the scientific literature - (e.g. NCBI gene/UniProt protein summaries, Gene Ontology annotations). Each modality can provide additive and complementary information when computing gene, and ultimately, cell representations.

**FIGURE 2.**
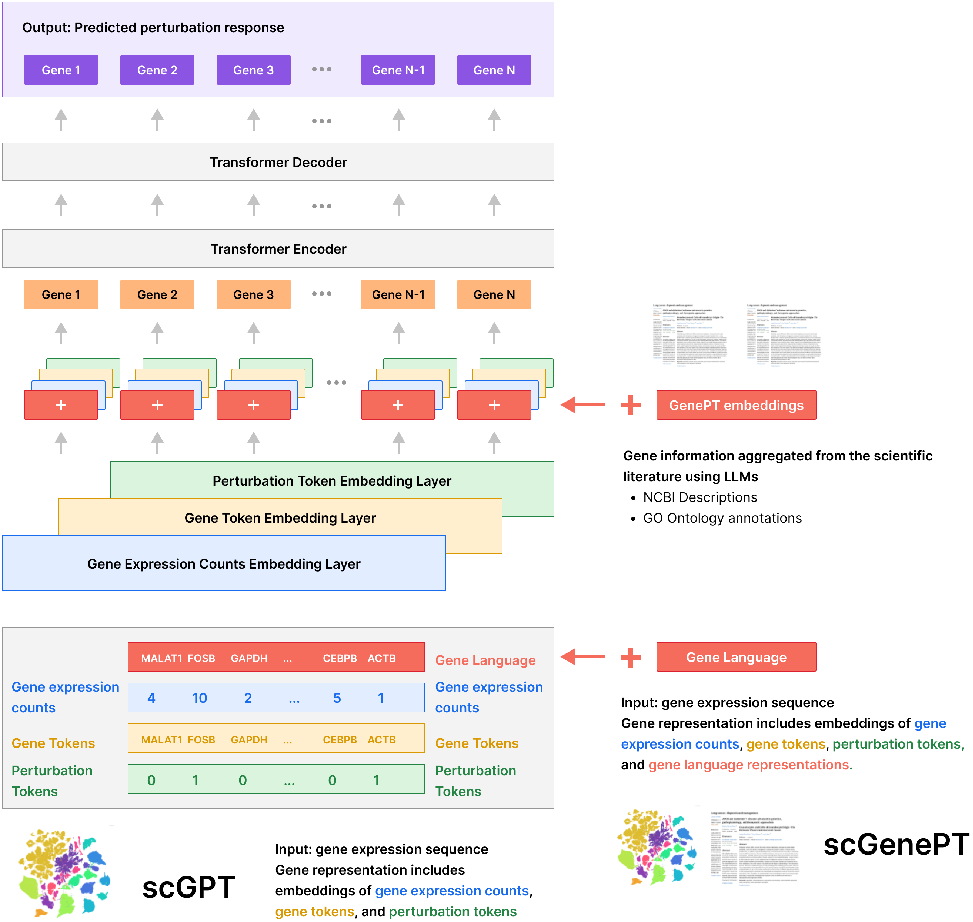
scGenePT Model Architecture. In scGPT, gene representations include gene expression counts, gene tokens and perturbation tokens. For scGenePT, each gene gets an additional representation, a gene language representation. Each of these different representations gets embedded using a separate embedding layer that gets learned during training. The gene embeddings are added element-wise to obtain one embedding per gene and then used as input to a Transformer Encoder layer. The outputs of this layer are decoded by a Transformer Decoder layer which generates predicted gene expression counts for each gene in the sequence. These are the predicted gene expression values after perturbation. The language embeddings are initialized from LLM-computed embeddings of gene information from the scientific literature, such as NCBI descriptions and Gene Ontology annotations.

**FIGURE 3.**
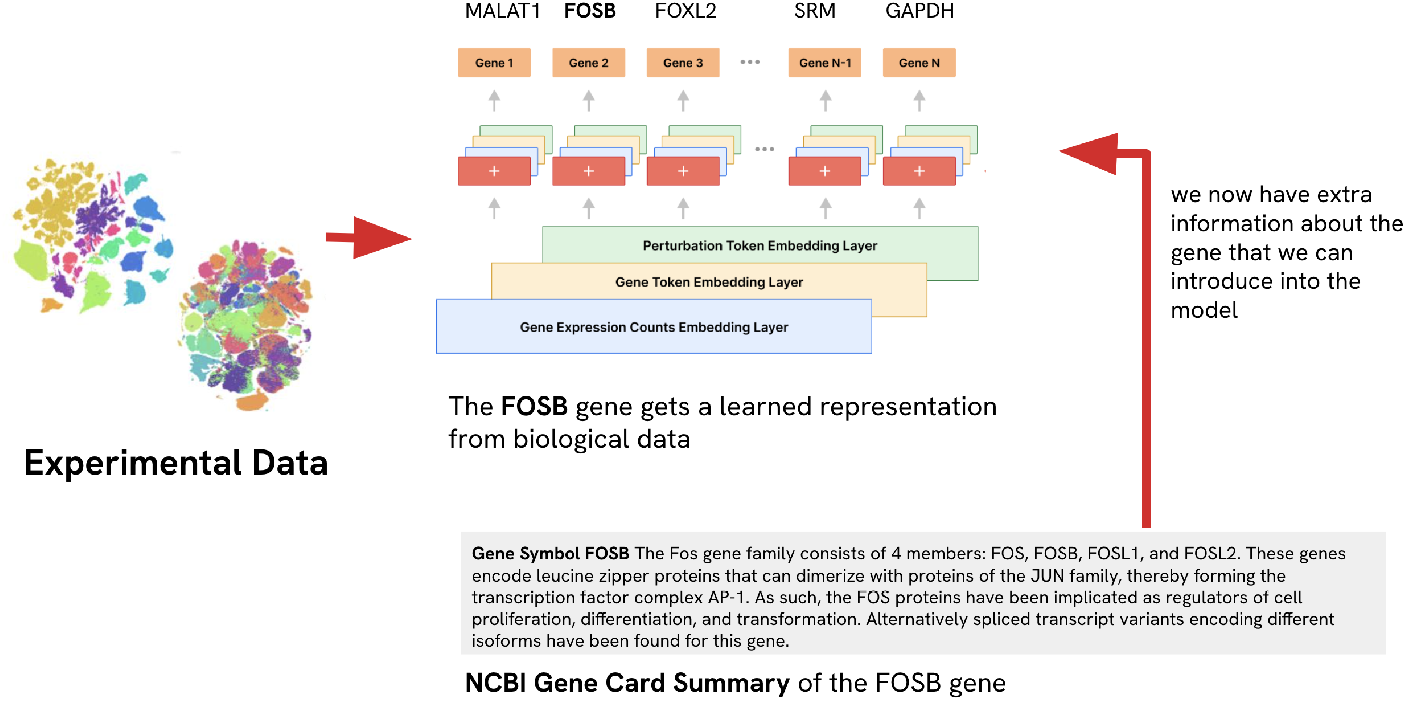
Example of introducing language gene representation into scGPT at the gene level

#### 2.2.1 The original scGPT Model Architecture

scGPT takes in a cell x gene matrix *X* ∈ ℝ^CxG^ containing *C* cells and *G* genes, where each entry *X*_*i,j*_ is the read count of an RNA molecule from scRNA-seq. Each gene is represented through a combination of its **gene token** (e.g. “FOSB”, “MALAT1”, etc), **gene counts** (e.g. 5, 10, 100, 0) and **perturbation conditions**. Gene tokens and perturbation conditions are learned through EmbeddingLayers, and gene counts are learned through an MLP. All of these different representations then get added element-wise to compute a final gene representation. A gene *g*^*i,j*^ in cell *c*^*i*^, gets an embedding *e*_*scGPT*_(*g*^*i,j*^) ∈ ℝ^d^:

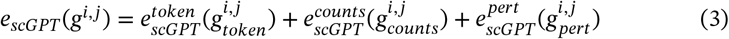

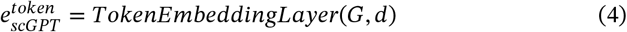

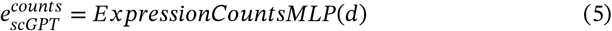

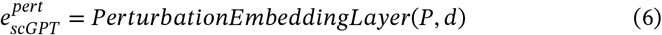

where **G** = number of genes, **P** = number of perturbation conditions and **d** = dimension of the learned gene embedddings

The perturbation condition *p*^*i,j*^ for genein *g*^*i,j*^ cell *c*^*i*^ is assigned as follows:

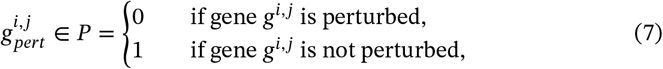

A cell observation *c*^*i*^ = [*g*^*i*,1^, *g*^*i*,2^, …*g*^*i,N*^] is then represented by a sampled sequence of *M* genes:

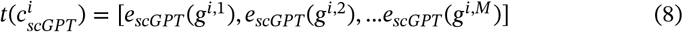

The cell representation 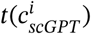 is fed into a TransformerEncoder, followed by a TransformerDecoder. The outputs of the TransformerDecoder layer are the predicted post-perturbation gene expression counts for each gene in the sequence.

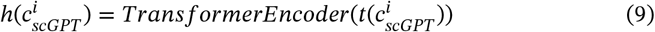

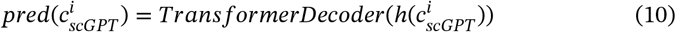

Note that for finetuning for pertution prediction, we initialize 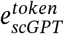 and 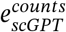 from the pre-trained model, while 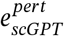 gets learned from scratch.

#### 2.2.2 Injecting language into scGPT

We retrieve textual information about each gene from different knowledge sources, such as NCBI gene card summaries [15], NCBI gene card summaries combined with UniProtKB [16] protein summaries for protein-coding genes, and gene annotations from the the Gene Ontology [17, 18], across three axes: molecular function, biological processes and cellular components. We do this for the list of all genes *G* that are in the scGPT vocabulary, doing an exact match on the gene names in the considered knowledge sources. For each knowledge source *s*, we use LLMs generated embeddings of the textual annotations. We hold the corresponding annotations in a matrix 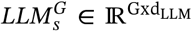 where *d*_*LLM*_ = dimensionality of the embedding model we use.

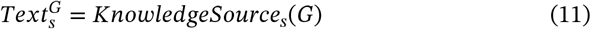

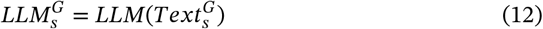

We retrieve the NCBI and NCBI + UniProtKB annotations from the genePT repository, and retrieve the gene annotations from the GO Ontology ourselves. We offer some examples of annotations and more details about the different types of knowledge sources in Table 1. Note that since the LLM embeddings are computed on a different representational space than the scGPT embeddings from Eq.(3), the two spaces need to be aligned. We do this using a linear projection layer. We then add the language embeddings as an additional representation at the gene level in Eq.(3). For a gene *g*^*i,j*^, the final representation becomes:

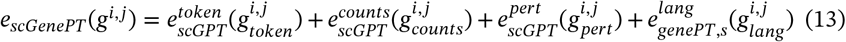

where

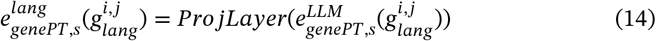

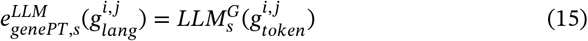

with 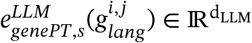 and 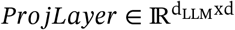

**TABLE 1.** Sources of scientific knowledge used for textual gene representation. Examples of textual gene annotations from each category for the gene FOSB.

| Knowledge Source | Example | Source/LLM |
| --- | --- | --- |
| NCBI Gene Card Summary | **Gene Symbol FOSB** The Fos gene family consists of 4 members: FOS, FOSB, FOSL1, and FOSL2. These genes encode leucine zipper proteins that can dimerize with proteins of the JUN family, thereby forming the transcription factor complex AP-1. As such, the FOS proteins have been implicated as regulators of cell proliferation, differentiation, and transformation. Alternatively spliced transcript variants encoding different isoforms have been found for this gene. | **genePT repository**<br>*GPT-3.5-text-embedding-ada-002* |
| NCBI Gene Card Summary + UniProtKB Protein Summary | **Gene Symbol FOSB** The Fos gene family consists of 4 members: FOS, FOSB, FOSL1, and FOSL2.[...] Alternatively spliced transcript variants encoding different isoforms have been found for this gene. **Protein summary:** Heterodimerizes with proteins of the JUN family to form an AP-1 transcription factor complex, thereby enhancing their DNA binding activity to gene promoters containing an AP-1 consensus sequence 5'-TGA[GC]TCA-3' and enhancing their transcriptional activity. ... *[truncated for brevity]* | **genePT repository/Computed**<br>*GPT-3.5-text-embedding-3-large/ada-002* |
| **Gene Ontology Annotation** - Molecular Function | **Molecular Function** terms: DNA-binding transcription factor activity, RNA polymerase II-specific, DNA-binding transcription activator activity, RNA polymerase II-specifify, DNA binding, protein binding, sequence-specific double-stranded DNA binding, RNA polymerase II cis-regulatory region sequence-specific DNA binding. **Annotation:** FOSB (enables) DNA-binding transcription factor activity | **Computed**<br>*GPT-3.5-text-embedding-ada-002* |
| **Gene Ontology Annotation** - Cellular Component | **Cellular Component** terms: nucleus, nucleoplasm, cytosol, intraceelular membrane-bounded organelle, chromatin. **Annotation:** FOSB (located in) nucleus | **Computed**<br>*GPT-3.5-text-embedding-ada-002* |
| **Gene Ontology Annotation** - Biological Process | **Biological Process** terms: negative regulation of transcription by RNA polymerase II, response to amphetamine, transcription by RNA polymerase II, female pregnancy. **Annotation:** FOSB (involved in) negative regulation of transcription by RNA polymerase II | **Computed**<br>*GPT-3.5-text-embedding-ada-002* |
| **Gene Ontology Annotation** - all - aggregation of all above | **Molecular Function:** DNA-binding transcription factor activity, RNA polymerase II-specific, DNA-binding transcription activator activity, ...; **Cellular Component:** nucleus, nucleoplasm, cytosol, intraceelular membrane-bounded organelle, chromatin; **Biological Process:** negative regulation of transcription by RNA polymerase II, response to amphetamine, transcription by RNA polymerase II, female pregnancy .. | **Computed**<br>*GPT-3.5-text-embedding-ada-002* |

We keep all of the following steps the same, except that we use layer normalization before feeding the summed embeddings into the TransformerEncoder module from Eq.(18). Through our experiments, we found that layer normalization helped in stabilizing the training when considering the two different embedding spaces and dimensionalities of the scGPT and LLMs embedding spaces. We offer more information on the different methods we tried in in the Appendix Section A.2.

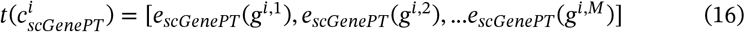

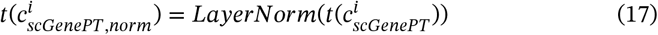

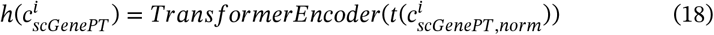

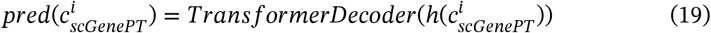

### 2.3 Source of Knowledge for Gene Language Representation

There are multiple sources of knowledge one can use to get textual representations for genes that can complement representations learned from experimental data. Essentially, there are many options for choosing the *KnowledgeSource*_*s*_ in Eq.11. We have explored using NCBI Gene Card Summaries, UniProt Protein Summaries and GO Ontology gene annotations. In the next sections, we describe each of these sources.

#### 2.3.1 NCBI Gene Card Summaries

NCBI Gene database (https://www.ncbi.nlm.nih.gov/gene/) [15] offers gene card summaries, which summarize gene function from the scientific literature. This can be viewed as an aggregation of scientific knowledge on a particular gene from the research articles published on it. We retrieve the embeddings from genePT [9], which computes them using the *GPT-3*.*5-text-embedding-ada-002* embedding model. We also perform ablation studies re-computing the embeddings using various LLAMA-3.1 model variants.

#### 2.3.2 UniProtKB

UniProtKB [16] offers protein summaries. For protein-coding genes, the protein summaries can be concatenated with the NCBI Gene Card information. Similarly to NCBI gene card summaries, we retrieve these embeddings from genePT and perform ablation studies re-computing the embeddings using LLAMA-3.1 model variants. Since these embeddings are computed using the *GPT-3*.*5-text-embedding-3-large* by genePT, we recompute them using the *GPT-3*.*5-text-embedding-ada-002* model, in order to make sure that the differences we see between different sources of textual representations are coming from the knowledge source itself rather than performance of embedding model choice.

#### 2.3.3 Gene Ontology (GO)

The Gene Ontology (http://geneontology.org) [17, 18] offers a computational representation of current scientific knowledge about the functions of genes. GO consists of three sub-ontologies with each providing annotations that describe one aspect of gene functionality: **molecular function, cellular component** and **biological process**.

- **GO-F: Molecular Function**: molecular level activities performed by gene products, such as “catalysis” or “transport”
- **GO-C: Cellular Component**: location of gene products relative to cellular compartments and structures, either in which (1) a gene product carries out a molecular function or (2) macromolecular complexes they are part of
- **GO-P: Biological Process**: the larger biological programs accomplished by multiple molecular activities (e.g. DNA Repair, signal transduction).

To these three categories that we obtain directly from the GO Ontology, we add a forth one, which is an aggregation of all three categories.

- **GO-all**: aggregation of GO-F (Molecular Function), GO-C (Cellular Component) and GO-P (Biological Process)

Examples of top annotations from each category can be seen in Figure 5. As shown in Table 1, each gene can have multiple annotations within each GO annotation category - for example, a gene can be tagged with multiple molecular functions, cellular components or biological processes. We’ve experimented with embedding each annotation separately and averaging the embeddings, and concatenating the multiple annotations together and embedding the concatenated string. For concatenation, we add a prefix specific to each knowledge source before concatenating all the annotations. We offer examples of how concatenation is performed from the raw annotations in Table 22.

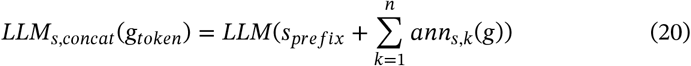

where

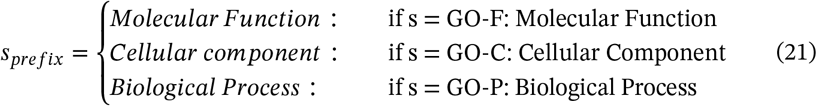

For averaging, we embed each annotation separately and then average the embeddings.

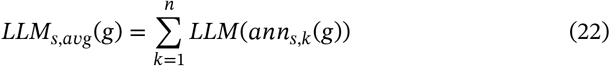

where *ann*_*s,k*_(*g*_*token*_) is the annotation for gene g with gene token *g*_*token*_ retrieved from knowledge sources

**FIGURE 4.**
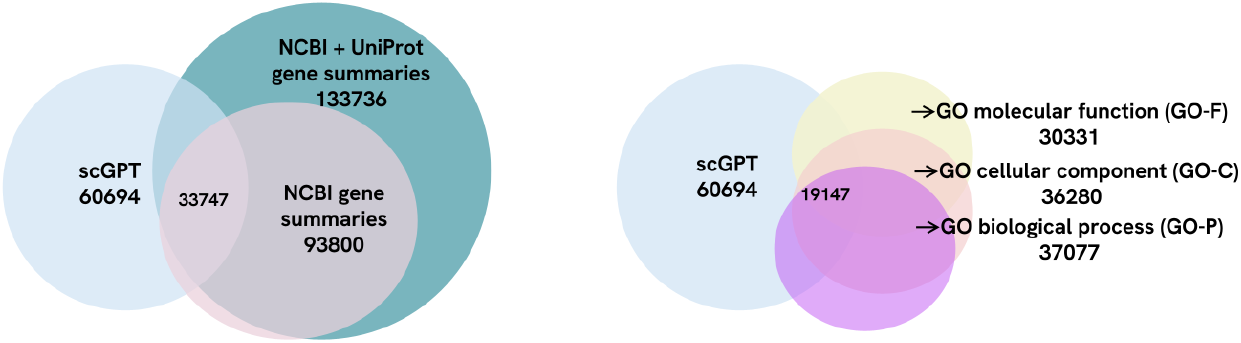
Number of gene embeddings for genes in the scGPT vocabulary across the considered knowledge sources. The intersection is represented in the middle.

**FIGURE 5.**
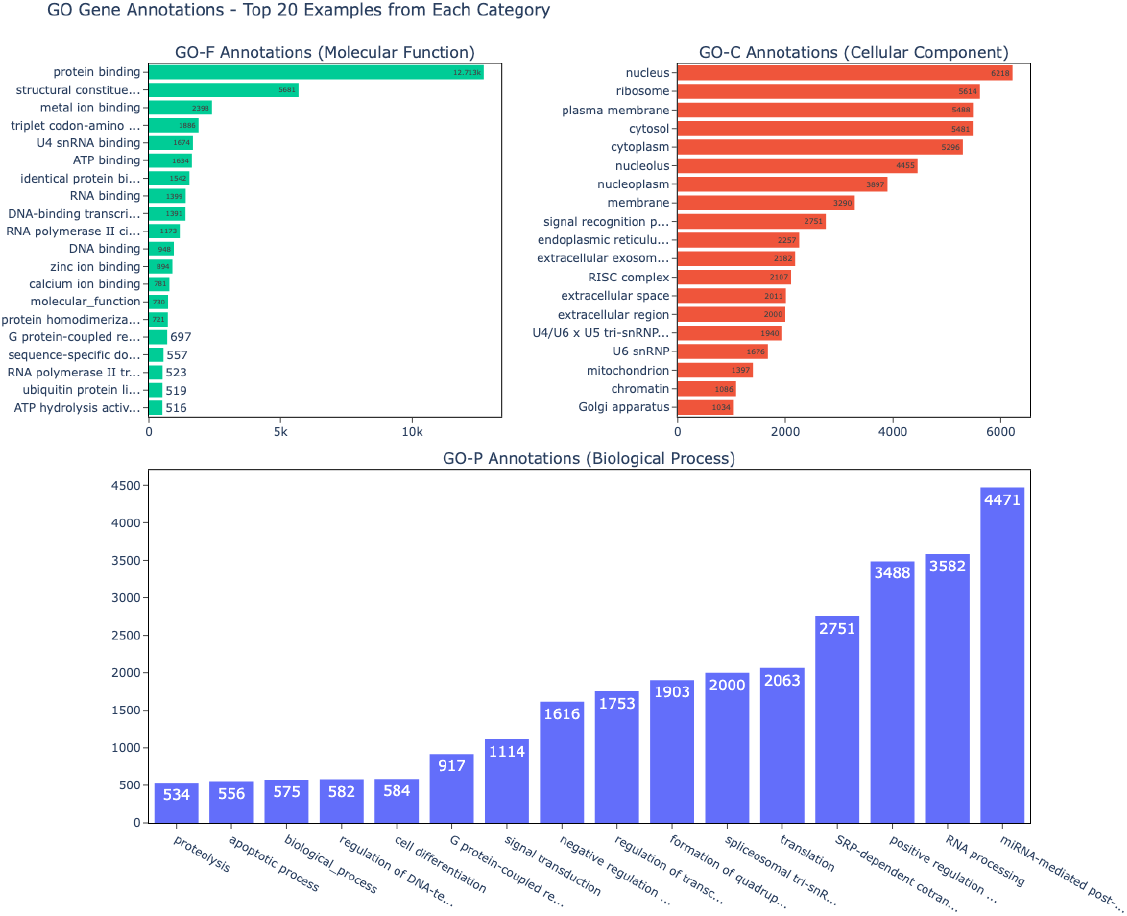
Distribution of GO Gene Annotations from each category. The top 20 terms from each category are displayed, together with the annotation counts over the GO Gene Ontology.

We’ve found slightly better results with concatenation, so we generally use the in *LLM*_*s,concat*_(*g*_*token*_)Equation 15 unless stated otherwise. We experimented with using both GPT-3.5, as well as different LLAMA-3.1 model variants as the embedding LLMs. In Figure 6 we can see a UMAP projection of the Gene Embeddings across the different sources. This projection uses Equation 20 for the GO annotations and the GPT-3.5 models for text embedding (*text-embedding-ada-002* for all annotations besides the NCBI + UniProt, which uses the *text-embedding-3-large*). We can see that while the NCBI and UniProt databases have a more balanced distribution of genes across the different gene functional classes, the GO annotations overwhelmingly contain protein coding genes. Similar plots are available in the Appendix for embeddings obtained using LLAMA-3.1-8b and LLAMA-3.1-70b models, as well as concatenation and average embedding mechanisms for the GO annotations. The gene functional classes have been obtained from the genePT GitHub repository https://github.com/yiqunchen/GenePT/blob/main/input_data/gene_info_table.csv.

**FIGURE 6.**
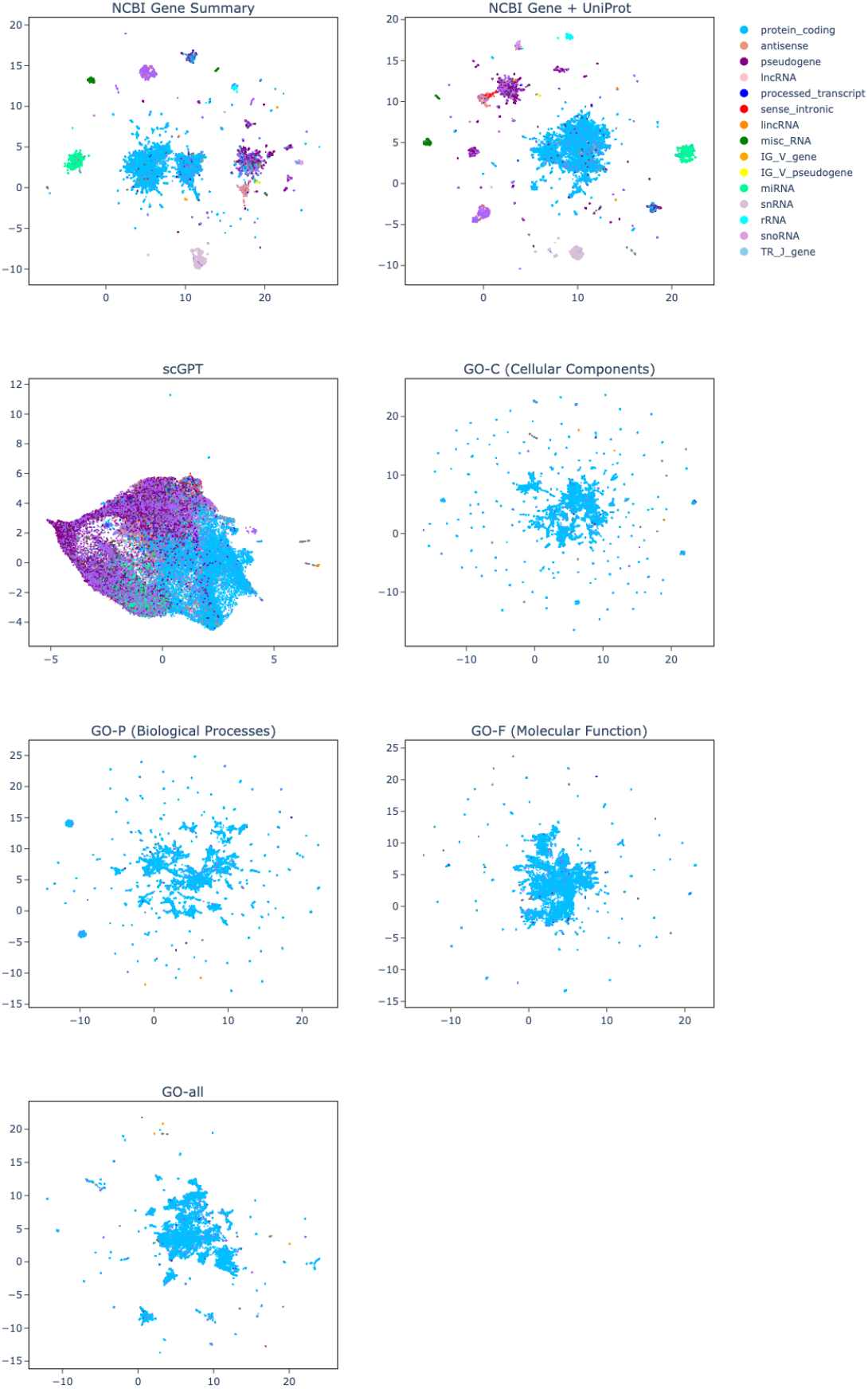
UMAP Projections of gene textual annotations embeddings, GPT-3.5, concatenating GO Annotations All annotations besides NCBI Gene + UniProt Summaries were embedded with GPT-3.5-ada embedding model. The NCBI Gene + UniProt Summaries was embedded with GPT-3.5-text-embedding-3-large model. The GO annotations used the concatenation method. Each color corresponds to a different gene functionality.

Specific examples of gene annotations from each source can be seen in Table 1, and the overlap between sources can be seen in Figure 4 and in Table 21.

## 3 Training

Models have been fine tuned from scGPT trained on the whole human corpus on GPU H100s. Each experiment has been run 5 different times, with 5 different seeds. We kept most of the hyperparameters the same as in scGPT, because we wanted to make the comparison as close as possible. Full list of parameters is in Appendix A.7. During training, one control sample is paired randomly with a perturbation and its response, which is considered ground truth. For each control/perturbed pair, we sample n = 1536 genes randomly and train on minimizing the MSE loss between ground truth and predicted perturbed response of the perturbation on control across all sampled genes. We keep the best model the one with the lowest MSE loss on the validation data.

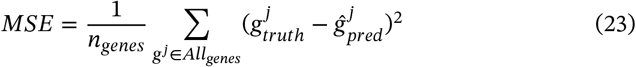

## 4 Evaluation

We evaluate our approach in single and two-gene perturbation settings. Single perturbation settings means that one gene gets perturbed, while in the two-gene setting, two genes get perturbed. In both cases, we are interested in predicting the effect of each type of perturbation on the gene expression profile of the cell. Notably, the perturbations of gene combinations are more challenging to predict, since they can have non-additive effects (e.g. the overall effect on the gene expression profile of having multiple genes perturbed at the same time is different than the cumulative effect of each gene perturbation).

### 4.1 Datasets

**Norman Dataset [20]** is a CRISPR perturb-seq dataset containing single and two-gene perturbations. We use a processed version of the dataset that contains 105 single and 131 two-gene combinations perturbations coming from 91k observations. Cells in the dataset are log-normalized and filtered to the top 5000 highly variable genes. The test is divided into train/val/test splits as shown in Figure 7. The test split is further post-processed in the following perturbation categories:

**FIGURE 7.**
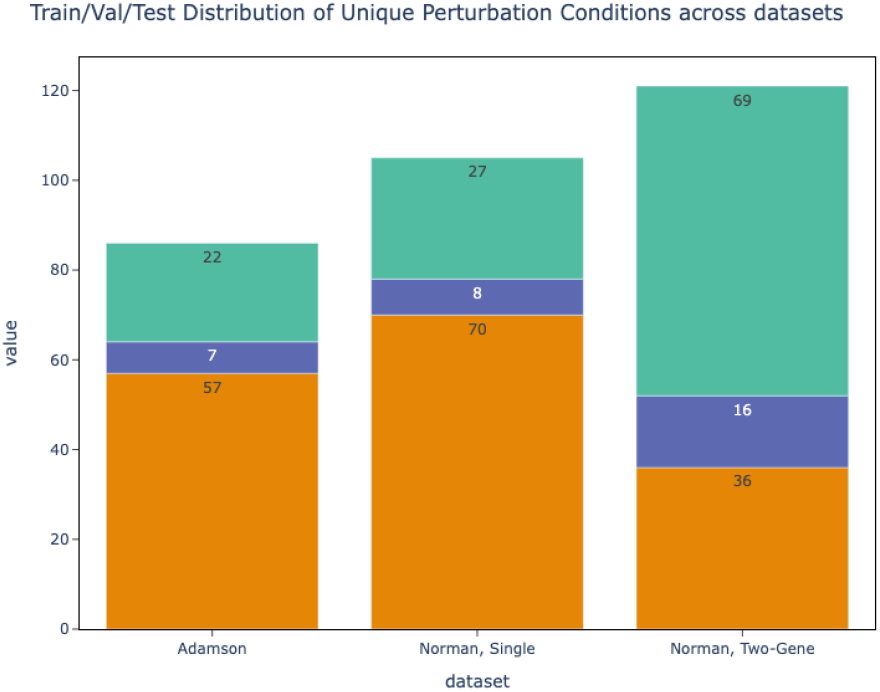
Train, Val, Test Splits Distribution of Unique perturbation Conditions across the datasets considered.

- **single** - one-gene perturbation, the gene has not been seen during training
- **two, seen 0/2** - two-gene perturbation, none of the genes has been seen perturbed during training
- **two, seen 1/2** - two-gene perturbation, one of the genes has been seen perturbed during training
- **two, seen 2/2** - two-gene perturbation, both genes has been seen perturbed during training

**Adamson Dataset [21]** is a 68k observations perturbation dataset containing 87 unique single perturbations using Perturb-seq. The data is log-normalized and filtered to the top 5000 highly variable genes.

For both Norman and Adamson datasets, we use the processed dataset versions from the GEARS package [14] v=0.0.2 and the corresponding train/val/test splits, which contain non-overlapping perturbations. The distribution of the data is seen in Figure 7 and in Table 2.

**TABLE 2.** Statistics of perturbation datasets used.

| Dataset | Types of per-<br>turbation | Number of<br>unique per-<br>turbations | Train/Val/Test<br>Splits | Number<br>of per-<br>turbated<br>observa-<br>tions | Number<br>of ctrl se-<br>quences | Total<br>number<br>of obser-<br>vations |
| --- | --- | --- | --- | --- | --- | --- |
| Adamson | single | 86 | 57/7/22 | 44340 | 24263 | 68603 |
| Norman | single | 105 | 70/8/27 | 48407 | 7353 | 91205 |
| Norman | two-gene | 131 | 36/16/69 | 35445 |  |  |

### 4.2 Metrics

For each perturbated sample in the test set, we sample a control sequence and predict the post-perturbation gene expression values from control, given the corresponding perturbation condition. Then, for each perturbation condition, we compute the mean true gene expression profile for that perturbation condition, as well as the mean predicted perturbation profile. We compute the metrics on the mean expresison values for each perturbation condition. We then average the metrics across **all** perturbation conditions. One perturbation condition corresponds to either one single-gene or two-gene perturbation condition.

Assuming we have P = *p*_1_, …*p*_*m*_ perturbation conditions, each with *p*_*i,k*_ samples, ∀*g*^*j*^ ∈ *G* we have:

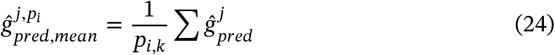

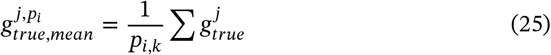

**MSE** We compute the **MSE** (Mean-Squared Error) between the mean true and predicted gene expression profiles after perturbation. We compute the metric on the whole set of genes, as well as on the top 20 differentially expressed genes, which we refer to as **MSE**_**Top20**_. We believe the latter, the MSE on the set of differentially expressed genes is a more signficant metric for model performance, given that the set of differentially expressed genes is the one that will have the most meaningful and substantial increase before and post-perturbation.

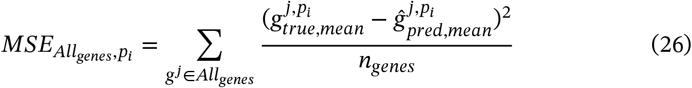

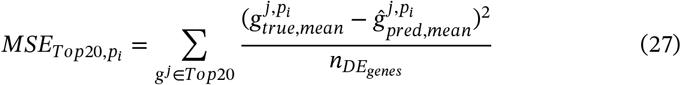

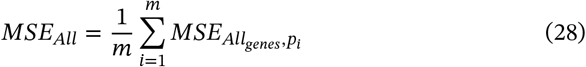

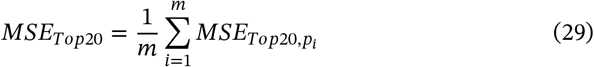

**Pearson Correlation** We take the **Pearson Correlation** between the mean true and mean predicted gene expression profile. We compute this metric for all genes, as well as the top 20 differrentially expressed genes, **Pearson Correlation**_**Top20**_.

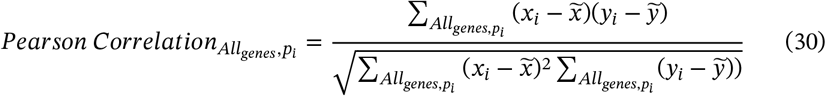

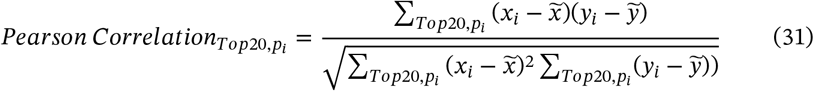

where 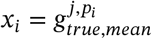 and 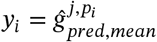

As with the MSE metrics, we take the average over all perturbation conditions:

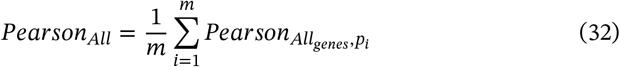

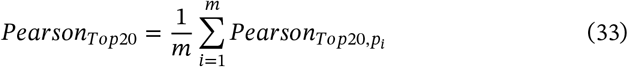

Similarly to recent literature [8, 22], we have also observed that the Pearson Correlation scores by themselves are not very meaningful and provide an inflated sense of model performance due to high correlation between non-essential genes in different cells. Hence, we don’t make heavy use of this metric in our analyses. Instead, we follow recent literature [8, 14, 22] that suggests using a variation of the metric that looks at the correlation between the gene expression value differences before and after perturbation as compared to control.

**Pearson CorrelationΔ** The **PearsonCorrelationΔ** looks at the correlation between the ground-truth and predicted post-perturbation expression values as compared to control. We compute the metric on the entire set of genes in the expression sequences, as well as the top 20 differentially expressed genes (including dropout genes), which we refer to as **PearsonCorrelationΔ**_**Top20**_. The main difference is that the values that we plug in into Equation 30 and Equation 31 are:

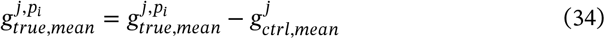

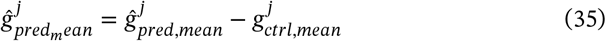

where 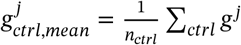

In our work, we have found that the metrics on the set of differentially expressed genes are more predictive than the metrics that take all the genes into consideration. This is consistent with recent literature [8]. Hence, while we compute all metrics, we take the **MSE**_**Top20**_ and **PearsonCorrelationΔ**_**Top20**_ to be more indicative of model

### 4.3 Baselines

**GEARS** We benchmark our models against GEARS [14], a state-of-the-art deep learning model for perturbation prediction that uses gene regulatory networks graphs to learn gene representations using graph-neural networks. We describe its architecture in more details in Related Work (6).

**Random Perturbation** We predict the post-perturbation gene expression vector of a perturbation taken at random from the training set of perturbations. Note that none of these perturbation conditions would be available during testing.

**Non-control-mean** Following Martens et al. [23] who found the non-control mean to be a strong baseline, we add this as a baseline in our analyses as well. In this setting, we take an average of all the perturbation responses from training data, and make that prediction for all the perturbations in the testing data.

## 5 Results

Throughout this section we use the naming convention **scGenePT_X** for scGenePT using Knowledge Source *X* for textual gene representation. The model correspondences are the following:

- **scGenePT**_**NCBI**_ = scGPT + NCBI Gene Card Summaries
- **scGenePT**_**NCBI+UniProt**_ = scGPT + NCBI Gene Card Summaries + UniProt Protein Summaries
- **scGenePT**_**GO**−**F**_ = scGPT + GO Molecular Functions Annotations
- **scGenePT**_**GO**−**C**_ = scGPT + GO Cellular Components Annotations
- **scGenePT**_**GO**−**P**_ = scGPT + GO Biological Processes Annotations
- **scGenePT**_**GO**−**all**_ = scGPT + GO_F + GO_C + GO_P

Metrics are reported over 5 different model runs, each ran with a different random seed. Error bars correspond to 95% intervals. The GO annotations are obtained through concatenation of individual annotation terms, as in Equation 20. The models use the *GPT-3*.*5-text-ada-002* embedding model unless stated otherwise. In each plot, we mark in bold the best model, and ^*^ the second best model.

### 5.1 Language provides additive value to biological representations in modeling single-cell perturbations

Our experiments show that adding textual representations for genes to the scGPT model architecture increases model performance in both the single-gene and two-gene perturbation settings across the Pearson Correlation and MSE metrics, both on the entire gene set and Top 20 DE genes. This can be seen in Table 3 for the two-gene perturbation Norman dataset and in Table 4 on the single-gene perturbation dataset Adamson. Notably, the increase is more proeminent in the two-gene perturbation setting, which is also more challenging due to potentially non-additive effects of gene interactions. In almost all cases, we observe an increase in performance regardless of the Knowledge Source used, which suggests that there is information carried by textual representations that augments the information carried in biological data.

**TABLE 3.** scGenePT Metrics on the Norman Dataset.

| Model | Pearson<br>Correlation<br>$\Delta_{\text{Top20}} \uparrow$ | Pearson<br>Correlation<br>$\Delta_{\text{all}} \uparrow$ | MSE<sub>Top20</sub> ↓ | MSE<sub>all</sub> ↓ |
| --- | --- | --- | --- | --- |
| scGPT | 0.665 ± 0.01 | 0.534 ± 0.02 | 0.223 ± 0.01 | 0.00421 ± 0.00 |
| scGenePT<sub>NCBI</sub> | 0.685 ± 0.03 | 0.548 ± 0.03 | 0.223 ± 0.03 | 0.00415 ± 0.00 |
| scGenePT<sub>NCBI+UniProt</sub> | **0.696 ± 0.01\*** | **0.557 ± 0.02** | **0.205 ± 0.02** | **0.00403 ± 0.00\*** |
| scGenePT<sub>GO-F</sub> | 0.686 ± 0.01 | **0.554 ± 0.02\*** | 0.216 ± 0.02 | 0.00405 ± 0.00 |
| scGenePT<sub>GO-C</sub> | 0.687 ± 0.03 | 0.550 ± 0.02 | 0.219 ± 0.01 | 0.00405 ± 0.00 |
| scGenePT<sub>GO-P</sub> | 0.682 ± 0.02 | 0.543 ± 0.02 | 0.220 ± 0.02 | 0.00412 ± 0.00 |
| scGenePT<sub>GO-all</sub> | **0.698 ± 0.02** | **0.554 ± 0.02\*** | **0.209 ± 0.02\*** | **0.00400 ± 0.00** |

**TABLE 4.** scGenePT Metrics on the Adamson Dataset.

| Model | Pearson<br>Correlation<br>$\Delta_{\text{Top20}} \uparrow$ | Pearson<br>Correlation<br>$\Delta_{\text{all}} \uparrow$ | MSE<sub>Top20</sub> ↓ | MSE<sub>all</sub> ↓ |
| --- | --- | --- | --- | --- |
| scGPT | 0.782 ± 0.02 | 0.589 ± 0.03 | 0.135 ± 0.01 | 0.00672 ± 0.00 |
| scGenePT<sub>NCBI</sub> | 0.779 ± 0.02 | 0.606 ± 0.03 | 0.133 ± 0.00 | 0.00654 ± 0.00 |
| scGenePT<sub>NCBI+UniProt</sub> | 0.784 ± 0.02 | **0.617 ± 0.03\*** | 0.129 ± 0.00 | **0.00620 ± 0.00** |
| scGenePT<sub>GO-F</sub> | 0.785 ± 0.02 | 0.611 ± 0.03 | 0.128 ± 0.00 | 0.00640 ± 0.00 |
| scGenePT<sub>GO-C</sub> | **0.791 ± 0.01** | **0.623 ± 0.02** | **0.125 ± 0.00** | **0.00622 ± 0.00** |
| scGenePT<sub>GO-P</sub> | **0.789 ± 0.01\*** | 0.609 ± 0.03 | **0.127 ± 0.00\*** | 0.00645 ± 0.00 |
| scGenePT<sub>GO-all</sub> | 0.787 ± 0.02 | 0.605 ± 0.03 | **0.127 ± 0.01\*** | 0.00641 ± 0.00 |

Note that the metrics presented in this section use concatenation of terms for GO Annotations as in Equation 20. We’ve also experimented with averaging the terms as in Equation 22 and found very little difference in performance. Hence, we’ve kept the concatenation method for all experiments and make the numbers for the averaging terms available in Appendix A.5.1 for the Adamson dataset and in Appendix A.5.2 for the Norman dataset.

The largest increase in performance in scGenePT compared to scGPT is in the two-gene perturbation dataset Norman. The two-gene perturbation settings are more challenging than the single-gene ones due to the potentially non-additive effects of gene interactions [20]. The dataset is split into multiple types of perturbations, which are of different levels of difficulty:

- **single gene** - one-gene perturbation, the gene has not been seen during training
- **two-gene, seen 0/2** - two-gene perturbation, neither of the genes has been seen perturbed during training
- **two-gene, seen 1/2** - two-gene perturbation, one of the genes has been seen perturbed during training
- **two-gene, seen 2/2** - two-gene perturbation, both genes have been seen perturbed during training

We see the biggest jump in model performance in the **two-gene, seen 0/2 setting**. This is remarkably also the hardest category to predict, due to the challenging nature of predicting non-additive effects of gene interactions, especially without the model having seen any of those genes perturbed during training. This tells us that representations aggregating information from the scientific literature are helping the model have a strong prior in predicting the effects of novel combinatorial perturbations. Hence, in the lack of experimental data, we can use available information from the literature as prior. While the biggest jump is in the **two-gene, seen 0/2 setting**, we also see an increase in performance across all four categories. Consequently, when experimental data exists, information from the literature can augment information learned from modeling biological data.

### 5.2 Different sources of scientific information have different performances in prediction

#### 5.2.1 For single gene perturbations, cellular component information provides the most added value

An interesting finding is that for single-gene perturbations, the GO Annotations Cellular Component (GO-C) information seems to be helping the most. This holds both on the Adamson dataset, as seen in Table 4, as well as in the single gene perturbation split of the Norman dataset, as seen in Table 5. In both of these cases, scGenePT_GO−C_ obtains the best metrics out of all the scGenePT_GO−X_ model variants. This tells us that subcellular localization is helpful in being able to predict effects of perturbation in single-gene perturbation settings. We have looked closely at two examples of single-gene perturbations: perturbing gene **POU3F2** and gene **CDKN1B**. In Figure 8 we can see a comparison of predictions generated by scGPT and scGenePT_GO−C_ for POU3F2. Both models have been finetuned on the Norman dataset, but POU3F2 has not been seen during training. We can see that the predictions generated by scGenePT_GO−C_ are more centered around the true ranges, capturing the directionality of the the predictions better. For example, gene FABP5, HSP90AB1, PRDX1, NPM1, TMSB10, PTMA are all better predicted as having a negative fold change over control by scGenePT_GO−C_, compared to scGPT which predicts a non-significant effect. According to NCBI Gene Card https://www.ncbi.nlm.nih.gov/gene/5454, overexpression of the protein encoded by POU3F2 is associated with an increase in the proliferation of melanoma cells. By adding the GO Cellular Component annotations, the model learns that this gene is localized mostly in: nucleoplasm, chromatin and transcription regulator complex. Localization of gene products in the cell plays an important role in their biological function, e.g. protein-protein interaction; regulation of gene expression, transportation of protein.

**TABLE 5.** **Pearson CorrelationΔ**_**Top20**_ Norman Dataset, Different Perturbation Categories.

| Model | single gene | two-gene,<br>seen 0/2 | two-gene,<br>seen 1/2 | two-gene,<br>seen 2/2 |
| --- | --- | --- | --- | --- |
| scGPT | $0.509 \pm 0.03$ | $0.579 \pm 0.08$ | $0.727 \pm 0.04$ | $0.844 \pm 0.03$ |
| scGenePT<sub>NCBI</sub> | $0.517 \pm 0.04$ | $0.649 \pm 0.03$ | $0.717 \pm 0.06$ | $0.857 \pm 0.04$ |
| scGenePT<sub>NCBI+UniProt</sub> | $0.507 \pm 0.03$ | **$0.660 \pm 0.03^*$** | **$0.745 \pm 0.03^*$** | **$0.872 \pm 0.03^*$** |
| scGenePT<sub>GO-F</sub> | **$0.521 \pm 0.05^*$** | $0.633 \pm 0.07$ | $0.731 \pm 0.01$ | $0.857 \pm 0.03$ |
| scGenePT<sub>GO-C</sub> | **$0.529 \pm 0.03$** | $0.615 \pm 0.1$ | **$0.746 \pm 0.02$** | $0.860 \pm 0.03$ |
| scGenePT<sub>GO-P</sub> | $0.507 \pm 0.03$ | $0.629 \pm 0.05$ | $0.736 \pm 0.02$ | $0.857 \pm 0.01$ |
| scGenePT<sub>GO-all</sub> | $0.510 \pm 0.04$ | **$0.662 \pm 0.04$** | **$0.745 \pm 0.00^*$** | **$0.874 \pm 0.03$** |

**FIGURE 8.**
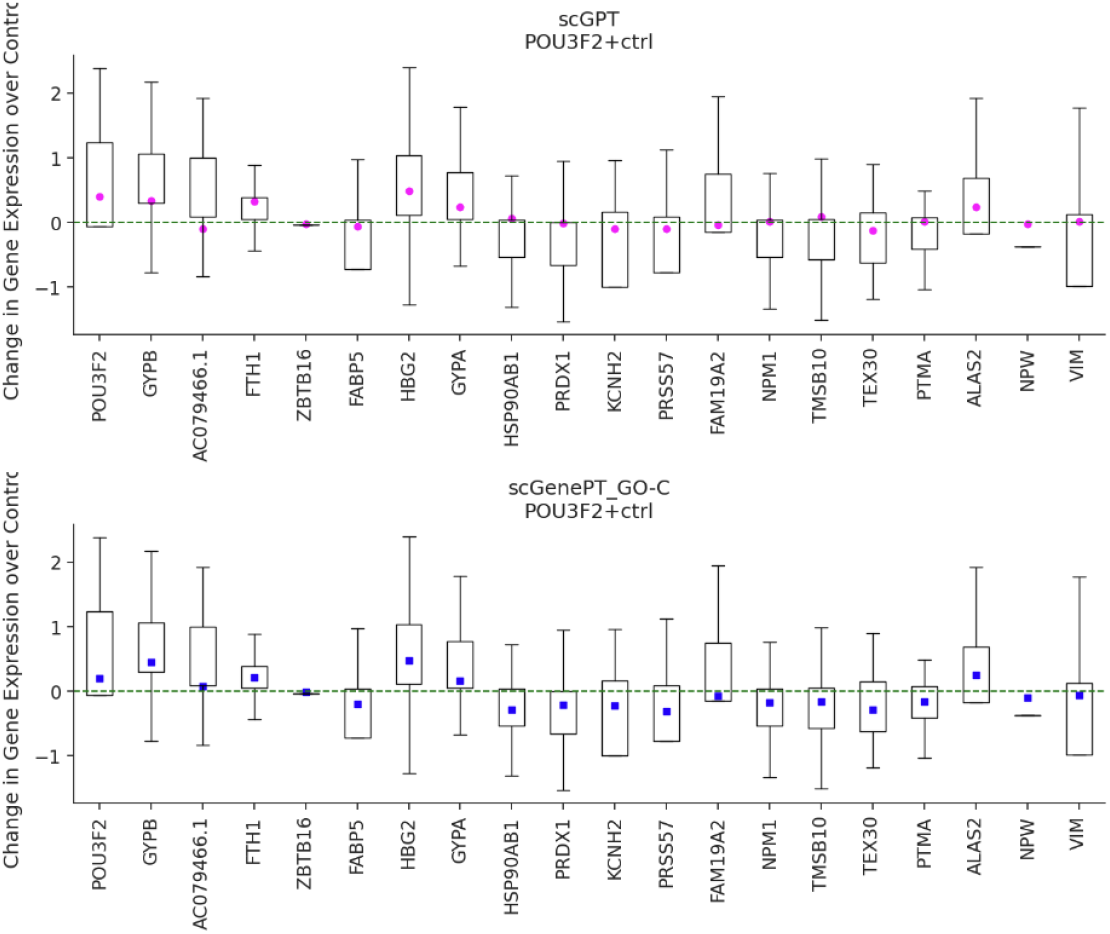
POU3F2 single-gene perturbation. Comparison of scGPT vs scGenePT predictions. Predictions are made over n = 300 randomly sampled controls.

In Figure 9, we can see a similar example for gene CDKN1B. Similarly, this gene has not been seen perturbed during training by the models used to generate the predictions. According to this gene’s NCBI Gene Card https://www.ncbi.nlm.nih.gov/gene/1027, mutations in this gene are associated with multiple enodcrine neoplasia type IV. We can see that scGenePT_GO−C_ predicts HSP90AA1, PTMA, RANBP1, CKS1B, PRDX1, PHF19 and NME1 as correctly down-regulated, as opposed to scGPT which predicts either neutral effect or positive fold change. In both cases, we speculate that the model learns to incorporate cellular location information to better predict gene expression change in response to genetic perturbation.

**FIGURE 9.**
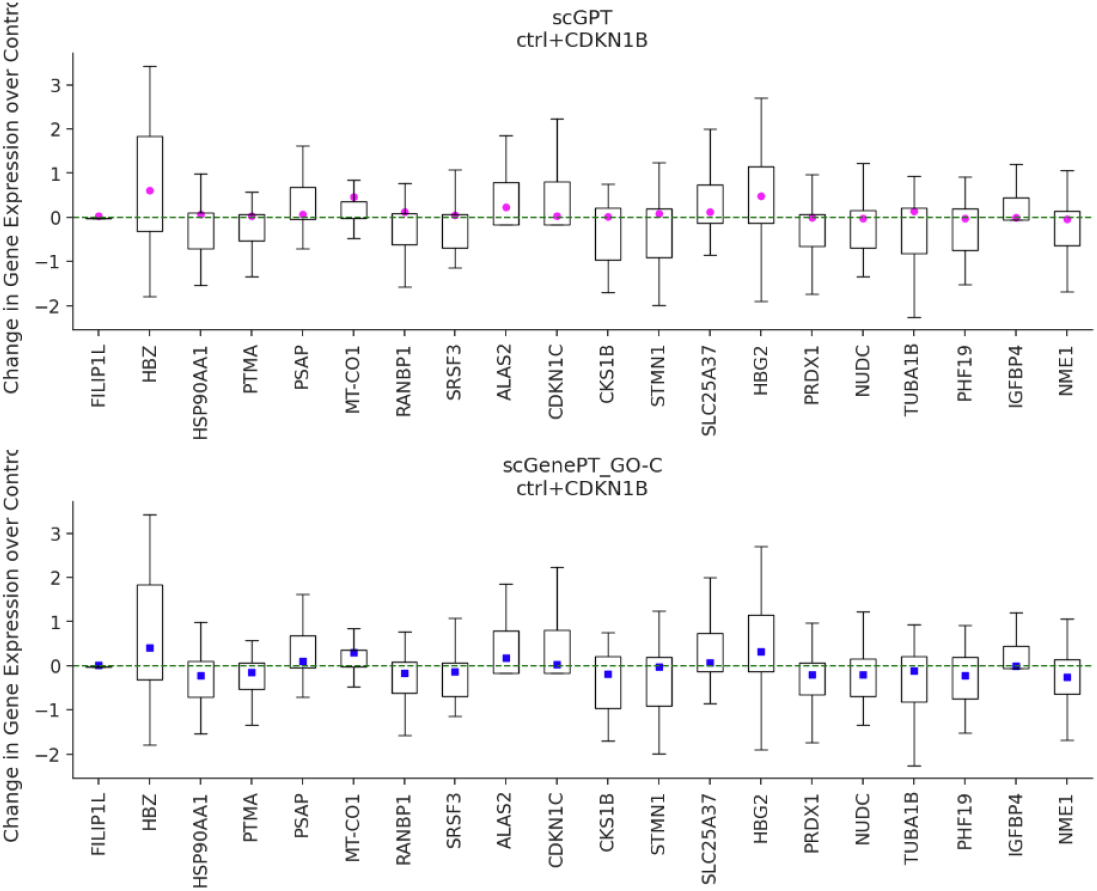
CDKN1B single-gene perturbation. Comparison of scGPT vs scGenePT predictions. Predictions are made over n = 300 randomly sampled controls.

**FIGURE 10.**
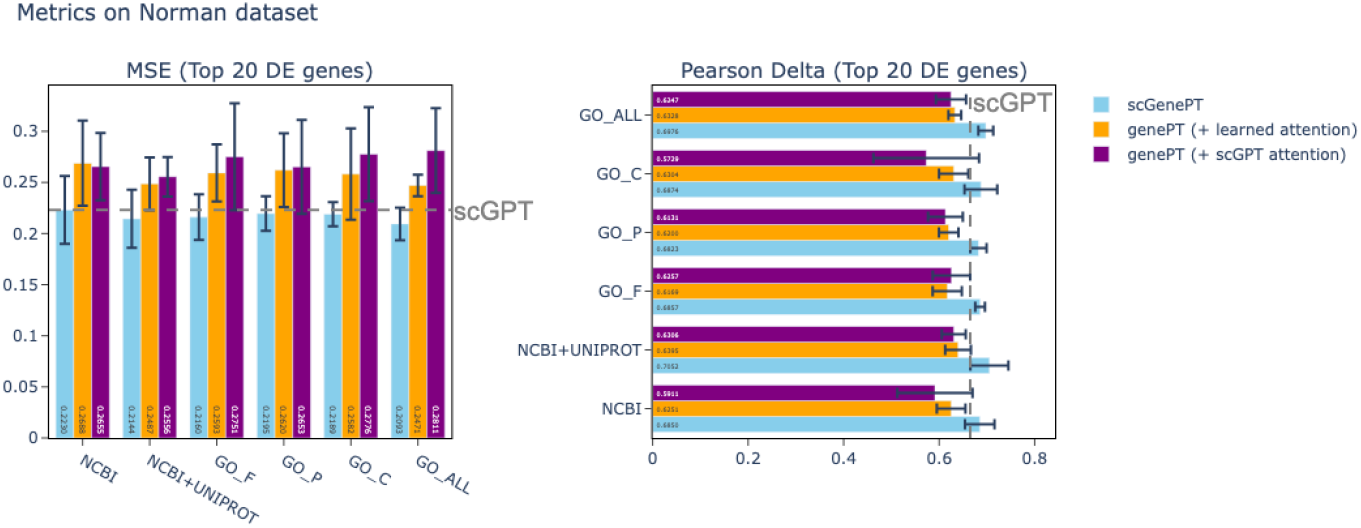
Comparison of performance of scGenePT with different language embeddings and language embeddings alone (genePT) on the test split of the Norman dataset. Note that we use the term **genePT** coined by [9] even when referring to our own embedded annotations

#### 5.2.2 For two-gene perturbations, protein summaries and aggregated annotations from the GO Ontology provide the most added value. The two hold similar information

As seen in Table3 and Table5, on the two-gene perturbation dataset Norman, the scGenePT_GO−all_ and scGenePT_NCBI+UniProt_ obtain the highest values, almost tied across all metrics. There are two things to note here:

- **scGenePT**_**NCBI+UniProt**_ **surpasses scGenePT**_**NCBI**_. Both are using embeddings computed with the same LLM (*GPT-3*.*5-text-embedding-ada-002*), and the only difference between the two is that scGenePT_NCBI+UniProt_ includes protein information. This tells us that for combinatorial perturbations, introducing not just gene information (from the NCBI Gene Card Summaries), but protein information for the protein-coding genes helps the model in predicting the effects of gene interactions on the transcriptome.
- **scGenePT**_**GO**−**all**_ **and scGenePT**_**NCBI+UniProt**_ **are tied almost across all metrics**. This shows that the two sources of information hold almost equivalent value - we can either aggregate all of the information across the GO Annotations landscape, spanning molecular function, biological process and cellular components, or take the information from the NCBI Gene Card Summaries combined with the protein information, and we would be encoding the same information.

The differences between model performance in the single vs two-gene perturbation settings might hint at the different levels of knowledge the model needs in different perturbation settings. One hypothesis would be that protein information becomes relevant in two-gene perturbation settings because of potential protein-protein interactions that don’t happen in the one-gene perturbation setting. Similarly, one could argue that in the one-gene perturbation setting, knowing the subcellular localization of the target becomes more straightforward information given there are no other potential non-additive effects of combinations of genes to consider. While more research would have to be done to explore these effects further, it is exciting to consider that we can create hypotheses on genes’ potential mechanism of action in different perturbation settings based on machine learning model performance alone.

#### 5.2.3 Different sources of information have complementary value

To probe further the complementarity aspect of these different knowledge sources, we ran ablation studies where we combined the NCBI+UniProt annotations with the different GO-annotations. On the single-gene perturbation dataset Adamson, the GO-F (molecular function), GO-P (biological processs) and GO-all (combination of GO-F + GO-P + GO-C) all benefit from combining the NCBI+UniProt and the GO annotations, scGenePT_NCBI+UniProt+GO−X_ surpassing the individual scGenePT _NCBI+UniProt_ and scGenePT_GO−X_ models. This can be observed in Table 8, Table 10 and Table 11.

**TABLE 6.** GO-C Annotations, POU3F2 and related genes.

| Gene | GO-C Cellular Component Annotation |
| --- | --- |
| POU3F2 | nucleoplasm, chromatin, transcription regulator complex |
| FABP5 | extracellular region, nucleus, nucleoplasm, cytoplasm, cytosol, plasma membrane, postsynaptic density, secretory granule membrane, azurophil granule lumen, synapse, extracellular exosome |
| HSP90AB1 | COP9 signalosome, protein folding chaperone complex, extracellular region, nucleus, nucleoplasm, cytoplasm, mitochondrion, cytosol, cell surface, membrane, secretory granule lumen, melanosome, neuronal cell body, dendritic growth cone, axonal growth cone, perinuclear region of cytoplasm, extracellular exosome, dynein axonemal particle, ficolin-1-rich granule lumen, protein-containing complex, aryl hydrocarbon receptor complex, HSP90-CDC37 chaperone complex, plasma membrane |
| PRDX1 | extracellular space, nucleus, cytoplasm, cytosol, melanosome, extracellular exosome |
| NPM1 | granular component, nucleus, nucleoplasm, nucleolus, cytoplasm, centrosome, cytosol, focal adhesion, membrane, spindle pole centrosome, protein-containing complex, protein-DNA complex, ribonucleoprotein complex |
| TMSB10 | cytoskeleton, cytoplasm |
| PTMA | nucleus, nucleoplasm, cytosol |

**TABLE 7.**
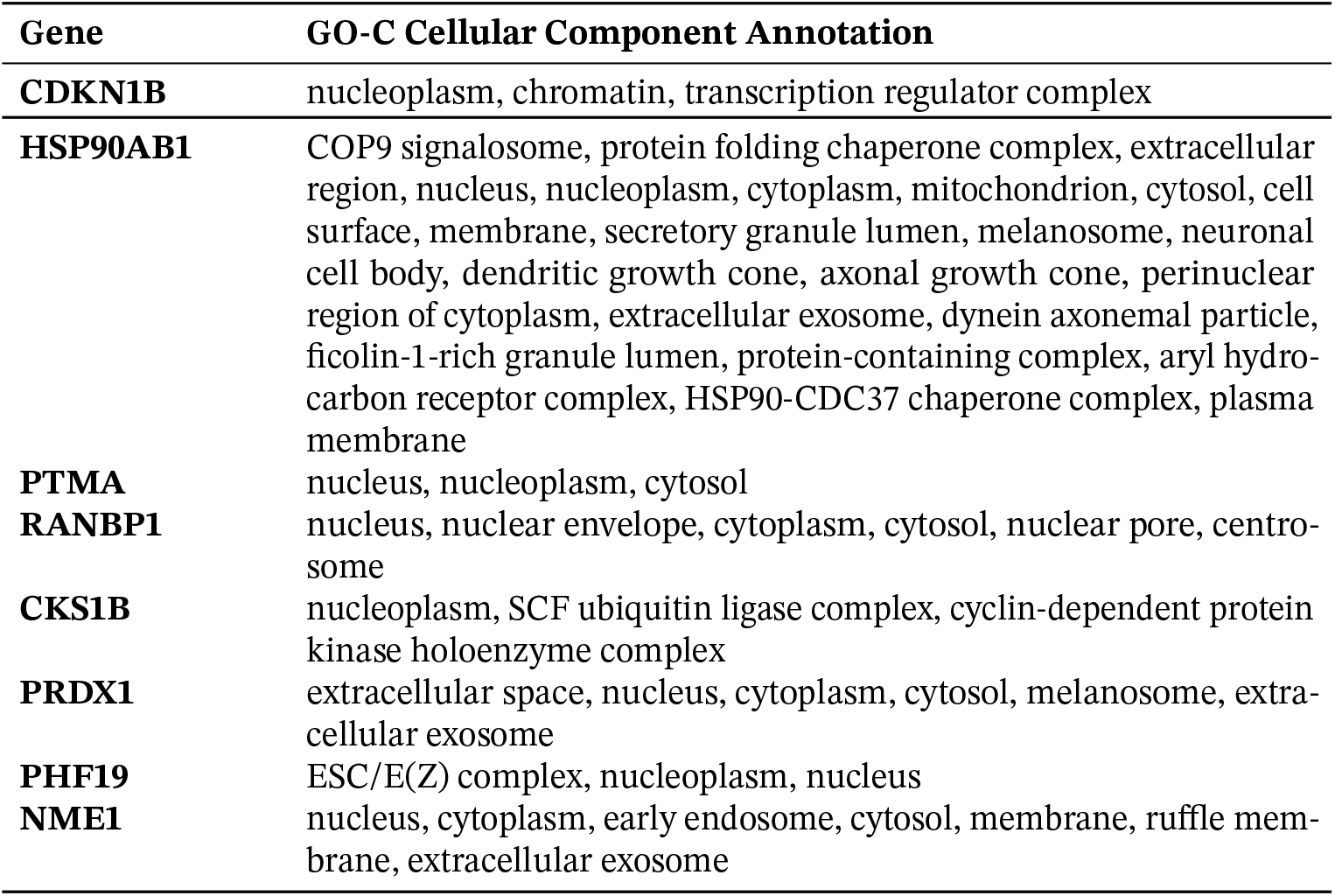
GO-C Annotations, CDKN1B and related genes.

**TABLE 8.**
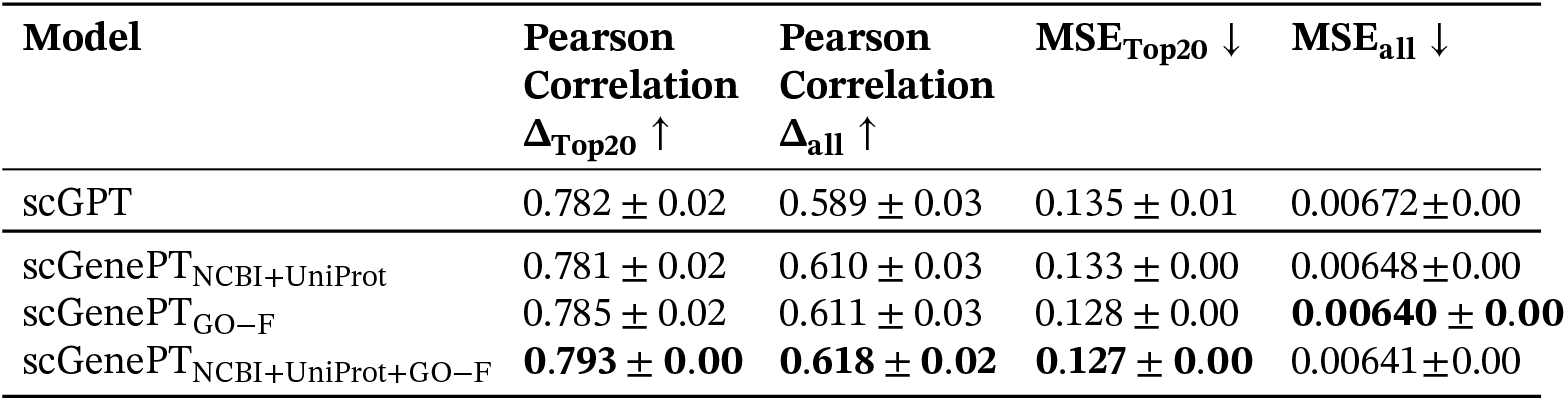
scGenePT (NCBI+UniProt + GO-F), Metrics on the Adamson Dataset.

**TABLE 9.** scGenePT (NCBI+UniProt + GO-C), Metrics on the Adamson Dataset.

| Model | Pearson<br>Correlation<br>$\Delta_{\text{Top20}} \uparrow$ | Pearson<br>Correlation<br>$\Delta_{\text{all}} \uparrow$ | MSE<sub>Top20</sub> ↓ | MSE<sub>all</sub> ↓ |
| --- | --- | --- | --- | --- |
| scGPT | 0.782 ± 0.02 | 0.589 ± 0.03 | 0.135 ± 0.01 | 0.00672±0.00 |
| scGenePT<sub>NCBI+UniProt</sub> | 0.781 ± 0.02 | 0.610 ± 0.03 | 0.133 ± 0.00 | 0.00648±0.00 |
| scGenePT<sub>GO-C</sub> | **0.791 ± 0.01** | **0.623 ± 0.02** | **0.125 ± 0.00** | **0.00622 ± 0.00** |
| scGenePT<sub>NCBI+UniProt+GO-C</sub> | 0.790 ± 0.01 | 0.617 ± 0.02 | 0.130 ± 0.00 | 0.00645±0.00 |

**TABLE 10.** scGenePT (NCBI+UniProt + GO-P), Metrics on the Adamson Dataset.

| Model | Pearson<br>Correlation<br>$\Delta_{\text{Top20}} \uparrow$ | Pearson<br>Correlation<br>($\Delta$)<sub>all</sub> ↑ | MSE<sub>Top20</sub> ↓ | MSE<sub>all</sub> ↓ |
| --- | --- | --- | --- | --- |
| scGPT | 0.782 ± 0.02 | 0.589 ± 0.03 | 0.135 ± 0.01 | 0.00672±0.00 |
| scGenePT<sub>NCBI+UniProt</sub> | 0.781 ± 0.02 | 0.610 ± 0.03 | 0.133 ± 0.00 | 0.00648±0.00 |
| scGenePT<sub>GO-P</sub> | 0.789 ± 0.01 | 0.609 ± 0.03 | **0.127 ± 0.03** | 0.00645±0.00 |
| scGenePT<sub>NCBI+UniProt+GO-P</sub> | **0.792 ± 0.00** | **0.620 ± 0.02** | **0.127 ± 0.00** | **0.00616 ± 0.00** |

**TABLE 11.** scGenePT (NCBI+UniProt+GO-all), Metrics on theAdamson Dataset.

| Model | Pearson<br>Correlation<br>$\Delta_{\text{Top20}} \uparrow$ | Pearson<br>Correlation<br>$\Delta_{\text{all}} \uparrow$ | MSE<sub>Top20</sub> ↓ | MSE<sub>all</sub> ↓ |
| --- | --- | --- | --- | --- |
| scGPT | 0.782 ± 0.02 | 0.589 ± 0.03 | 0.135 ± 0.01 | 0.00672±0.00 |
| scGenePT<sub>NCBI+UniProt</sub> | 0.781 ± 0.02 | 0.610 ± 0.03 | 0.133 ± 0.00 | 0.00648±0.00 |
| scGenePT<sub>GO-all</sub> | 0.787 ± 0.02 | 0.605 ± 0.03 | 0.127 ± 0.01 | 0.00641±0.00 |
| scGenePT<sub>NCBI+UniProt+GO-all</sub> | **0.793 ± 0.00** | **0.617 ± 0.00** | **0.126 ± 0.00** | **0.00627 ± 0.00** |

However, the effect does not hold GO-C annotations, where combining the annotations leads to worse performance than each of the individual models, as shown in Table 9. This tells us that the GO-P and GO-F annotations might have complementary value to the NCBI+UniProt annotations, whereas the GO-C annotations do not. Note that for the metrics presented in this section, NCBI+UniProt annotations have been embedded with *GPT-3*.*5-text-embedding-3-large* and all of the GO annnotations have been embedded with *GPT-3*.*5-text-embedding-ada-002*.

### 5.3 Language serves as a good prior for modeling single-cell perturbation, but is not sufficient

If language alone could replace biologically-learned representations, then models trained on textual gene representations alone should achieve the same performance as models that learn those biological representations from expression data during scGPT pre-training, given that the model architecture is the same. To test this hypothesis, we kept the scGPT model architecture, but removed the biologically-learned representations (gene tokens, gene counts and learned attention) from scGPT pre-training.

We call these model variations **genePT**_**X**_, where similarly as before, **X** represents the specific knowledge source used to retrieve the language gene embeddings. The results in Table 12 and Table 14 on Norman show that across the board, given any knowledge source **X**, models trained on language data alone do not reach the performance of scGPT, and consequently, scGenePT.

**TABLE 12.** genePT vs scGenePT, Norman Dataset, **Pearson Corr Δ**_**Top20**_.

| Model | Pearson Corr $\Delta_{\text{Top20}} \uparrow$ | Model | Pearson Corr $\Delta_{\text{Top20}} \uparrow$ (learned attention) | Pearson Corr $\Delta_{\text{Top20}} \uparrow$ (scGPT attention) |
| --- | --- | --- | --- | --- |
| scGPT | $0.665 \pm 0.01$ | scGPT | $0.665 \pm 0.01$ | $0.665 \pm 0.01$ |
| scGenePT<sub>NCBI</sub> | **$0.685 \pm 0.03$** | genePT<sub>NCBI</sub> | $0.625 \pm 0.03^*$ | $0.591 \pm 0.08$ |
| scGenePT<sub>NCBI+UniProt</sub> | **$0.696 \pm 0.01$** | genePT<sub>NCBI+UniProt</sub> | $0.640 \pm 0.03^*$ | $0.631 \pm 0.03$ |
| scGenePT<sub>GO-F</sub> | **$0.686 \pm 0.01$** | genePT<sub>GO-F</sub> | $0.617 \pm 0.03$ | $0.626 \pm 0.04^*$ |
| scGenePT<sub>GO-C</sub> | **$0.687 \pm 0.03$** | genePT<sub>GO-C</sub> | $0.630 \pm 0.03^*$ | $0.573 \pm 0.11$ |
| scGenePT<sub>GO-P</sub> | **$0.682 \pm 0.02$** | genePT<sub>GO-P</sub> | $0.620 \pm 0.02^*$ | $0.613 \pm 0.04$ |
| scGenePT<sub>GO-all</sub> | **$0.698 \pm 0.02$** | genePT<sub>GO-all</sub> | $0.633 \pm 0.01^*$ | $0.625 \pm 0.03$ |

**TABLE 13.** genePT vs scGenePT, Norman Dataset, **Pearson Corr Δ**_**All**_.

| **Model** | **Pearson Corr ($\Delta_{All}$) ↑** | **Model** | **Pearson Corr ($\Delta_{All}$) ↑**<br>**(learned attention)** | **Pearson Corr ($\Delta_{All}$) ↑**<br>**(scGPT attention)** |
| --- | --- | --- | --- | --- |
| scGPT | $0.534 \pm 0.03$ | scGPT | $0.534 \pm 0.03$ | $0.534 \pm 0.03$ |
| scGenePT<sub>NCBI</sub> | **$0.548 \pm 0.03$** | genePT<sub>NCBI</sub> | $0.302 \pm 0.02$ | $0.318 \pm 0.02^*$ |
| scGenePT<sub>NCBI+UniProt</sub> | **$0.557 \pm 0.02$** | genePT<sub>NCBI+UniProt</sub> | $0.322 \pm 0.02^*$ | $0.320 \pm 0.02$ |
| scGenePT<sub>GO-F</sub> | **$0.554 \pm 0.02$** | genePT<sub>GO-F</sub> | $0.307 \pm 0.03$ | $0.313 \pm 0.03^*$ |
| scGenePT<sub>GO-C</sub> | **$0.550 \pm 0.02$** | genePT<sub>GO-C</sub> | $0.312 \pm 0.02$ | $0.315 \pm 0.02^*$ |
| scGenePT<sub>GO-P</sub> | **$0.543 \pm 0.02$** | genePT<sub>GO-P</sub> | $0.309 \pm 0.01$ | $0.329 \pm 0.02^*$ |
| scGenePT<sub>GO-all</sub> | **$0.554 \pm 0.02$** | genePT<sub>GO-all</sub> | $0.321 \pm 0.02^*$ | $0.305 \pm 0.01$ |

**TABLE 14.** genePT vs scGenePT, Norman Dataset. **MSE**_**Top20**_.

| **Model** | **MSE<sub>Top20</sub> ↓** | **Model** | **MSE<sub>Top20</sub> ↓**<br>**(learned attention)** | **MSE<sub>Top20</sub> ↓**<br>**(scGPT attention)** |
| --- | --- | --- | --- | --- |
| scGPT | $0.223 \pm 0.02$ | scGPT | $0.223 \pm 0.02$ | $0.223 \pm 0.02$ |
| scGenePT<sub>NCBI</sub> | **$0.223 \pm 0.03$** | genePT<sub>NCBI</sub> | $0.269 \pm 0.04$ | $0.266 \pm 0.03^*$ |
| scGenePT<sub>NCBI+UniProt</sub> | **$0.214 \pm 0.03$** | genePT<sub>NCBI+UniProt</sub> | $0.249 \pm 0.03^*$ | $0.256 \pm 0.02$ |
| scGenePT<sub>GO-F</sub> | **$0.216 \pm 0.02$** | genePT<sub>GO-F</sub> | $0.259 \pm 0.03^*$ | $0.275 \pm 0.05$ |
| scGenePT<sub>GO-C</sub> | **$0.219 \pm 0.01$** | genePT<sub>GO-C</sub> | $0.258 \pm 0.05^*$ | $0.278 \pm 0.05$ |
| scGenePT<sub>GO-P</sub> | **$0.220 \pm 0.02$** | genePT<sub>GO-P</sub> | $0.262 \pm 0.04^*$ | $0.265 \pm 0.05$ |
| scGenePT<sub>GO-all</sub> | **$0.209 \pm 0.02$** | genePT<sub>GO-all</sub> | $0.247 \pm 0.01^*$ | $0.281 \pm 0.04$ |

Even after adding the attention mechanism to the genePT models, models are still lagging behind. Hence, the learned token and counts embeddings during pre-training are essential to the model performance coming from experimental data. Metrics on the entire set of genes, as well as on the top 20 differentially expressed genes are consistent. Metrics for Adamson are available in the Appendix Table 27 and Table 29 and show a similar trend.

### 5.4 Language can help biologically-informed models in surpassing other model architectures that have specific biological knowledge hard-coded in model architecture

In our analyses, we notice that scGenePT models often surpass GEARS in the single-gene perturbation setting - as seen on the Adamson dataset in Table 18 and in the single-gene setting of the Norman dataset in Table 17. However, the same does not always hold in the two-gene perturbation setting as seen in Table 16. In fact, looking at the detailed results in Table 17, we see that the biggest increase in the two-gene setting comes from GEARS performing well on the **two-gene, seen 0/2** setting. This is the category where the model has to predict the effect of two-gene perturbations where none of the genes has been seen during training. Note that this is also the setting in which the language representations add the most value to the scGPT models. When there is information about at least one of the gene perturbed, such as in the **two-gene, seen 1/2** and **two-gene, seen 2/2** settings, scGenePT model variants often surpass GEARS. Considering that GEARS is learning embeddings directly from the gene regulatory network graphs, this can reinforce the value of structured information as priors to guide model training in situations where training data is limited or non-existent. Given that predicting the effect of a random perturbation from the training set, or the non-ctrl-mean obtains values similar to GEARS, it can also point to a dataset-specific issue. We offer a more detailed discussion on this in the Discussion 8 section.

**TABLE 15.** genePT vs scGenePT, Norman Dataset. **MSE**_**All**_.

| **Model** | **MSE<sub>All</sub> ↓** | **Model** | **MSE<sub>All</sub> ↓**<br>**(learned attention)** | **MSE<sub>All</sub> ↓**<br>**(scGPT attention)** |
| --- | --- | --- | --- | --- |
| scGPT | $0.00421 \pm 0.00$ | scGPT | $0.00421 \pm 0.00$ | $0.00421 \pm 0.00$ |
| scGenePT<sub>NCBI</sub> | **$0.00415 \pm 0.00$** | genePT<sub>NCBI</sub> | $0.00904 \pm 0.00$ | $0.00896 \pm 0.00^*$ |
| scGenePT<sub>NCBI+UniProt</sub> | **$0.00403 \pm 0.00$** | genePT<sub>NCBI+UniProt</sub> | $0.00874 \pm 0.00^*$ | $0.00880 \pm 0.00$ |
| scGenePT<sub>GO-F</sub> | **$0.00405 \pm 0.00$** | genePT<sub>GO-F</sub> | $0.00900 \pm 0.00^*$ | $0.00903 \pm 0.00$ |
| scGenePT<sub>GO-C</sub> | **$0.00405 \pm 0.00$** | genePT<sub>GO-C</sub> | $0.00892 \pm 0.00^*$ | $0.00903 \pm 0.00$ |
| scGenePT<sub>GO-P</sub> | **$0.00412 \pm 0.00$** | genePT<sub>GO-P</sub> | $0.00899 \pm 0.00$ | $0.00890 \pm 0.00^*$ |
| scGenePT<sub>GO-all</sub> | **$0.00407 \pm 0.00$** | genePT<sub>GO-all</sub> | $0.00885 \pm 0.00^*$ | $0.00904 \pm 0.00$ |

**TABLE 16.** scGenePT Metrics on the Norman Dataset.

| Model | Pearson<br>Correlation<br>$\Delta_{\text{Top20}} \uparrow$ | Pearson<br>Correlation<br>$\Delta_{\text{all}} \uparrow$ | MSE<sub>Top20</sub> ↓ | MSE<sub>all</sub> ↓ |
| --- | --- | --- | --- | --- |
| scGPT | 0.665 ± 0.01 | 0.534 ± 0.02 | 0.223 ± 0.01 | 0.00421 ± 0.00 |
| scGenePT<sub>NCBI+UniProt</sub> | 0.696 ± 0.01 | 0.557 ± 0.02 | **0.205 ± 0.02** | **0.00403 ± 0.00** |
| scGenePT<sub>GO-all</sub> | **0.698 ± 0.02** | 0.554 ± 0.02* | 0.209 ± 0.02* | **0.00400 ± 0.00** |
| GEARS | **0.710 ± 0.00** | **0.573 ± 0.00** | **0.177 ± 0.00** | 0.00510 ± 0.00 |
| random perturbation | 0.604 ± 0.03 | 0.495 ± 0.00 | 0.340 ± 0.00 | 0.00488 ± 0.00 |
| non-ctrl-mean | 0.608 ± 0.00 | **0.571 ± 0.00\*** | 0.337 ± 0.00 | 0.00466 ± 0.00 |

**TABLE 17.** **Pearson Correlation Δ**_**Top20**_ Norman Dataset, Different Perturbation Categories.

| Model | single gene | two-gene,<br>seen 0/2 | two-gene,<br>seen 1/2 | two-gene,<br>seen 2/2 |
| --- | --- | --- | --- | --- |
| scGPT | $0.509 \pm 0.03$ | $0.579 \pm 0.08$ | $0.727 \pm 0.04$ | $0.844 \pm 0.03$ |
| scGenePT<sub>NCBI+UniProt</sub> | $0.507 \pm 0.03$ | $0.660 \pm 0.03^*$ | **$0.745 \pm 0.03^*$** | **$0.872 \pm 0.03^*$** |
| scGenePT<sub>GO-C</sub> | **$0.529 \pm 0.03$** | $0.615 \pm 0.1$ | **$0.746 \pm 0.02$** | $0.860 \pm 0.03$ |
| scGenePT<sub>GO-all</sub> | $0.510 \pm 0.04$ | $0.662 \pm 0.04$ | **$0.745 \pm 0.00^*$** | **$0.874 \pm 0.03$** |
| GEARS | $0.448 \pm 0.01$ | **$0.785 \pm 0.00$** | $0.737 \pm 0.01$ | $0.869 \pm 0.00$ |
| random perturbation | $0.397 \pm 0.04$ | **$0.771 \pm 0.06$** | $0.630 \pm 0.01$ | $0.619 \pm 0.00$ |
| non-ctrl-mean | $0.397 \pm 0.00$ | $0.741 \pm 0.00$ | $0.656 \pm 0.00$ | $0.640 \pm 0.00$ |

**TABLE 18.** scGenePT Metrics on the Adamson Dataset.

| Model | Pearson<br>Correlation<br>$\Delta_{\text{Top20}} \uparrow$ | Pearson<br>Correlation<br>$\Delta_{\text{all}} \uparrow$ | MSE<sub>Top20</sub> ↓ | MSE<sub>all</sub> ↓ |
| --- | --- | --- | --- | --- |
| scGPT | 0.782 ± 0.02 | 0.589 ± 0.03 | 0.135 ± 0.01 | 0.00672 ± 0.00 |
| scGenePT<sub>GO-C</sub> | **0.791 ± 0.01** | 0.623 ± 0.02 | **0.125 ± 0.00** | 0.00622 ± 0.00 |
| GEARS | 0.730 ± 0.00 | 0.691 ± 0.00 | 0.157 ± 0.00 | 0.00516 ± 0.00 |
| random perturbation | 0.726 ± 0.03 | **0.693 ± 0.00** | 0.151 ± 0.00 | **0.00482 ± 0.00** |
| non-ctrl-mean | 0.729 ± 0.00 | **0.709 ± 0.00\*** | 0.149 ± 0.00 | **0.00456 ± 0.00\*** |

### 5.5 Ablation studies

**Effect of different biological components** Our analyses show that language can augment representations learned from experimental data. In order to test what aspect of the biologically-learned representations from experimental data is the most useful, we tested separate elements. We found that taking either the tokens or the counts representations out of scGPT model performance leads to the model not learning any meaningful representations. Taking the tokens out had a more significant effect. Hence, the token and count representations are both essential for capturing the variety in experimental data. This holds in both single-gene in Table 20 and two-gene perturbation settings in Table 19.

**TABLE 19.** Ablation Studies, Norman Dataset.

| Model | Pearson Corr<br>$\Delta_{\text{Top20}} \uparrow$ | MSE<sub>Top20</sub> ↓ | Pearson Corr<br>$\Delta_{\text{All}} \uparrow$ | MSE<sub>All</sub> ↓ |
| --- | --- | --- | --- | --- |
| scGPT | $0.665 \pm 0.01$ | $0.223 \pm 0.02$ | $0.534 \pm 0.02$ | $0.0042 \pm 0.00$ |
| scGPT (-counts) | $0.035 \pm 0.00$ | $2.784 \pm 0.02$ | $0.128 \pm 0.00$ | $0.1619 \pm 0.00$ |
| scGPT (-tokens) | $0.143 \pm 0.01$ | $0.546 \pm 0.02$ | $0.137 \pm 0.00$ | $0.0115 \pm 0.00$ |

**TABLE 20.** Ablation Studies, Adamson Dataset.

| Model | Pearson Corr<br>$\Delta_{\text{Top20}} \uparrow$ | MSE<sub>Top20</sub> ↓ | Pearson Corr<br>$\Delta_{\text{All}} \uparrow$ | MSE<sub>All</sub> ↓ |
| --- | --- | --- | --- | --- |
| scGPT | $0.782 \pm 0.02$ | $0.135 \pm 0.01$ | $0.589 \pm 0.03$ | $0.0067 \pm 0.00$ |
| scGPT (-counts) | $0.257 \pm 0.00$ | $7.776 \pm 0.04$ | $0.044 \pm 0.00$ | $0.2834 \pm 0.00$ |
| scGPT (-tokens) | $0.023 \pm 0.05$ | $0.323 \pm 0.03$ | $-0.092 \pm 0.02$ | $0.0271 \pm 0.00$ |

## 6 Related Work

### 6.1 Large language models for single-cell biology

There have been a number of foundation transformer-based models developed for single-cell biology. These models generally use semi-supervised learning to train on unlabeled data. They are then fine tuned on specialized tasks, such as cell type annotation, batch integration or perturbation prediction. Models in this category include scGPT [8], GeneFormer [11], UCE [10] and scMulan[13], which are trained on large-scale scRNAseq data from single-cell atlases and repositories and use experimental data (e.g. scRNAseq counts) as a modality to learn from. On the other hand, genePT [9] is a foundation model for single-cell biology that uses language as a modality to represent genes through LLM-computed embeddings of NCBI descriptions. In our work, we extend the scGPT model architecture to include genePT embeddings and experiment with different sources of knowledge besides the ones considered by genePT. Our model makes use of and extends both the scGPT and genePT model architectures and shows the additive and complementary value of the representations learned by these two models. Moreover, while most of these foundation models are trained on one modality, our model incorporates both scRNAseq counts and language training on two modalities at the same time.

### 6.2 Models incorporating biological experimental data and language

The main research question we explore in this paper is the value added by incorporating language as a modality into representations learned from experimental data. Specifically for perturbation prediction in single-cell data, the idea of using LLM-computed gene embeddings has been explored by Martens et al. [23] who showed that text and protein embeddings are complementary and additive when used as priors in Gaussian Process models and that combining the two surpasses GEARS in predicting perturbation prediction on the one-gene perturbation Replogle dataset [24]. We do not test on the Replogle dataset in our work, but we consider it as a next step, and believe that comparing the performance of the different approaches would be interesting to make. While not evaluated specifically on modeling perturbation, some models for single-cell biology incorporate joint training on single-cell and language. CellWhisperer [25] uses a CLIP framework to learn a joint embedding space between transcriptomes obtained from the GEO database, encoded using GeneFormer [11], and textual annotations encoded using BioBERT [26], exploring the idea of interrogation of cells with natural language. ChatNT [27] also uses separate encoders for natural language and DNA sequences, learning a projection layer to align the two spaces. Similarly in our work, we learn a projection layer to map from language-based embeddings to biologically learned embeddings of scGPT. However, the scope of both CellWhisperer and ChatNT is different, focusing on answering questions through natural language, whereas we treat language as a modality for encoding prior information from the scientific literature. Cell2Sentence [28] and CeLLAma [29] build cell sentences by transforming gene expression profiles into sentences in natural language, treating single-cell tasks through a framework that is entirely based on natural language. These models have not been tested on perturbation prediction, and do not explicitly use language as a way to encode information from the scientific literature as a prior.

### 6.3 Models for perturbation prediction

Many of the models mentioned so far fall into the category of *foundation models* (scGPT, UCE, GeneFormer, genePT) or *instruction tuning models* (ChatNT, Cell2Sentence). In this paradigm, perturbation is a downstream fine-tuning task. There are a number of model architectures that have been designed exclusively for perturbation prediction. GEARS [14] is a knowledge-graph based deep learning model that incorporates prior knowledge from gene-gene relationships to predict transcriptional responses to single- and multi-gene perturbations. Our model introduces prior knowledge from the scientific literature into an existing foundation model (scGPT). While GEARS introduces prior information as a knowledge graph representation, we introduce it through language as aggregation from the literature. Hence, differences in performance between the two approaches can also be indicative of the value of different types of representations for biological data - graphs versus language. Like with GEARS, we are also able to predict the effects of unseen single- and multi-gene perturbations. While this is possible due to the nature of the scGPT model architecture, we show in our analyses that language can increase performance. CPA [30] is a generative model built for single-cell perturbation prediction that disentangles cellular states into basal and perturbation latent variables. SAMS-VAE [22] is a joint generative model that introduces a sparse additive mechanism in order to disentangle perturbed samples to identify perturbation-specific latent subspaces. While we don’t compare against generative models for perturbation in this work, we believe that the idea of introducing language information as priors into generative models is an exciting area of future research.

## 7 Code and Data Availability

### Code

The scGPT model architecture has been retrieved from the scGPT Github repository https://github.com/bowang-lab/scGPT in July 2024. Its modified version to incorporate the gene embeddings, as well as all model code used for training and inference, code used for analyses and demonstration notebooks will be made available upon publication, together with trained model checkpoints. The scGPT model has been finetuned from scGPT whole human checkpoint available at https://drive.google.com/drive/folders/1oWh_-ZRdhtoGQ2Fw24HP41FgLoomVo-y.

### Data

The genePT embeddings (NCBI gene card summaries embedded with *gpt-3*.*5-text-embedding-ada* and NCBI gene card + UniProtKB protein summaries embedded with *gpt-3*.*5-text-3-large*) were retrieved from https://zenodo.org/records/10833191. We recomputed the NCBI gene card + UniProtKBprotein summaries with *gpt-3*.*5-text-embedding-ada* to make the ablations between different knowledge sources more fair. These embeddings, together with the embeddings of gene GO functional terms annotations, obtained with GPT-3.5, as well as LLAMA-3.1-8b and LLAMA-3.1-70b will be made available upon publication. The gene annotations from the GO ontology were retrieved in 2024-07-15 from 10.5281/zenodo.10536401.

For the Adamson and Norman splits, we use the dataloaders and splits from GEARS [14], version 0.0.2. All other packages and versions will be made available on the code repository.

## 8 Discussion

### Effect of language in augmenting experimental data

In our analyses, we found that introducing textual representations can complement biologically learned representations from experimental data. One interesting finding of our work is that different types of information seem to help in different ways. In particular, for single-gene perturbation settings, subcellular localization retrieved through the Cellular Components gene annotations from the GO Gene Ontology information seems to be helping the most. This hints to the fact that in the single-gene perturbation setting, where there are no gene interaction effects from perturbed genes, knowing the subcellular location of where genes bind is more useful information for the model rather than other types of information. In the two-gene perturbation setting, the UniProt protein summaries seem to help the most. This can be seen as achieving similar generalization to hard-coding prior biological knowledge into model architecture through protein-protein interaction networks, similarly to models like GEARS, by attending on the initial textual descriptions describing the effects of the protein-coding genes. We are encouraged that a transcriptomic foundation model like scGPT can learn to generalize in structured fashion implicitly through attention and language embeddings without explicitly needing access to structures like graphs (e.g. protein-protein interaction networks, gene regulatory networks). The exciting aspect of our work is that we observe a significant increase in generalization without requiring architecture changes for explicit wiring of prior knowledge, additional pretraining, or inductive biases, while achieving generalization akin to or better than models with explicit inductive biases. This suggests that our current understanding about the performance ceiling of transcriptomic foundation models is incomplete and incorporation of curated knowledge beyond experimental data may provide signficant gains.

### Structured information is useful as prior in predicting effects of novel perturbation interactions

Generally in the single-gene perturbation case, scGPT and scGenePT model variants surpass models like GEARS that are using information from biological knowledge graphs to learn gene representations. The place where GEARS still outperforms transformer-based deep learning models is in predicting novel two-gene perturbations where none of the genes has been seen during training - the **two-gene, seen 0/2** setting. This is also the place where the textual gene representations help the most in increasing scGPT performance. This could hint to the importance of using priors from structured knowledge sources to inform prediction of novel perturbation interactions. However, predicting the effect of a random perturbation from the training set or the non-ctrl-mean obtain similar values to GEARS. This can tell us that there might be a dataset-specific issue where the effects of this particular set of genes in the 0 seen of 2 combination are tightly related to the the effect of the perturbations in the training set. Lastly, our goal is to study the effect of language in complementing experimental data, using scGPT as our test model architecture. This means that we are inheriting model performance from scGPT, and it is possible that this is simply a case where GEARS performs better. More analyses would have to be done here in order to understand the effect in this particular setting. However, the fact that there are instances where we beat or match models such as GEARS that use rich specific inductive biases, such as regulatory networks without encoding this information in the model architecture itself speaks to the tremendous potential that large deep learning models have in learning these deep biological features embedded in structured data during training.

### Aligning disjoint embedding spaces

In order to incorporate the textual gene embeddings into the scGPT model architecture, the textual embeddings need to be aligned to the scGPT embedding space. This is non-trivial, not just because these two models have different dimensions, but also because they have been trained on different data distributions. We have explored multiple ways of aligning the two, which we describe in Appendix A.2. Testing our multiple approaches on the Norman dataset, we have found that the ways in which the embeddings are aligned can have a considerable effect on downstream performance. Exploring the optimal way of aligning biological and textual representation spaces is an exciting area that needs to be explored more. For the purpose of the experiments we carried out in this work, we kept a configuration that gave reasonable results for both scGPT and scGenePT, so as to not bias the alignment choice towards either model. However, we believe that more exploration could be done in this area that could lead to better performance.

### Metrics and Limitations

One of the biggest limitations for the task of perturbation prediction remains evaluation [8]. The most commonly used metrics are MSE (or MAE), the Pearson Correlation Δ and the Average Treatment Effect as proposed by Bereket et al. in [22], either on the entire set of genes or on the Top-K differentially expressed genes. In our analyses, we used MSE and Pearson Correlation Δ, giving more weight to these metrics on the set of Top-20 differentially expressed genes. We found that there are cases where model variants that obtain the best metrics on the entire set of genes do not perform similarly on the differentially expressed genes, and vice versa. We chose to focus on the set of differentially expressed genes because these are the genes that would change the most given a perturbagen, so are more informative from a biological perspective. Moreover, we noted that often times the results that obtained the highest metrics on the entire set of genes were random perturbation or non-ctrl-mean, which means that it is possible that these metrics would be optimizing set of genes that are constant to a baseline cell profile, rather than specific to a perturbation effect. Similarly, there wasn’t always a correlation between the models obtaining the smallest MSE and the highest Pearson Correlation values, as one would expect. These contradictory results make it hard to evaluate models fairly and with certainty. While we report metrics commonly used in the field and believe that the signal justifies our results, we want to acknowledge that perturbation modeling evaluation will need to improve as pertaining to overall dataset availability and evaluation procedures to infer more generalizable insights. We are excited about the ongoing work in the field of perturbation benchmarking and will continue re-evaluating our methods as the field evolves.

### Using pre-trained models

We also want to highlight the potential limitation of our results given our strategy to modify and base this work off a specific pre-trained model. Of course scGenePT inherits performance characteristics from its core model, and it would be interesting future work to evaluate the outcome of training from scratch with these features in mind or adjusting the modeling choices of the underlying transcriptomic foundation model to increase performance.

## 9 Conclusion

In our work, we explored the value of adding language as an additional modality to augment representations learned from experimental data. We did this by injecting textual embeddings into scGPT, a popular foundation model for single-cell data. We experimented with different knowledge sources for gene representations, such as NCBI gene descriptions, NCBI gene descriptions combined with UniProt protein summaries, and GO Annotations, spanning molecular function, biological processes and cellular components. We found that adding language embeddings improves performance of biologically learned representations, and that different sources have different effects. Subcellular localization through GO-C Cellular Components Annotations provides the highest value for single-gene perturbations and UniProt protein summaries are the most useful for predicting two-gene perturbation interaction effects. We have learned that, when used properly, **language can be used to complement biological representations for single-cell data**. We saw that language embeddings by themselves don’t reach the performance of biologically learned representations, showing that **biology and language are two complementary representations, but language is not sufficient on its own**. We’ve showed that we are able to match or beat models that have extra knowledge embedded into the model architecture directly (such as GEARS) by using information about genes from the literature or databases as a covariate. This suggests that we have not exhausted the potential of black box ML models like transformers to increase in performance within the data regimes relevant to bespoke biology tasks, but may need to look into careful curation outside the experimental data to afford the models better generalization. Overall, our work showcases the value of adding language information from the literature to augment representations learned from experimental data. We believe there are a lot of exciting opportunities to build on top of this work and probe harder into specific sources of knowledge and ways of embedding them in deep learning models. We are excited about how this line of work can inform building multi-modal models based on language and experimental data, and about using language in meaningful ways to inform, guide and complement biological experiments.

## 10 Acknowledgment

We would like to thank Michaela Torkar for manuscript revisions and Yasmine Mabene for help with retrieving the GO Gene Annotations.

## A Appendix

### A.1 Intersection of genes across pairs of knowledge sources considered

**TABLE 21.** Intersection of genes across pairs of knowledge sources considered.

| Gene Knowledge Source 1 | Gene Knowledge Source 2 | # common genes |
| --- | --- | --- |
| **scGPT** n=60694 genes | **NCBI** n=93800 genes | 33747 |
|  | **NCBI+UniProt** n=133736 genes | 33752 |
|  | **GO-F** n=30331 genes | 17865 |
|  | **GO-C** n=36280 genes | 18492 |
|  | **GO-P** n=37077 genes | 17482 |
|  | **GO-all** n=44301 genes | 19147 |
| **NCBI** n=93800 genes | **NCBI+UniProt** n=93800 genes | 93800 |
| **GO-C** n=36280 genes | **GO-P** n=93800 genes | 29604 |
| **GO-C** n=36280 genes | **GO-F** n=93800 genes | 26683 |
| **GO-F** n=30331 genes | **GO-P** n=93800 genes | 23408 |

### A.2 Aligning embeddings

We have explored multiple ways of aligning the textual gene embeddings to the scGPT embedding space:

- Linear Projection: Linear projection that maps from the genePT embedding to the scGPT embedding space during scGPT finetuning
  – w/o emb normalization: no embedding normalization
  – with embedding normalization
- Linear Projection + Batch normalization: Linear_projection + batch normalization before feeding the added embeddings into transformer
- Linear Projection + Layer normalization: Linear_projection + layer normalization before feeding the added embeddings into transformer
- Linear Projection + Dropout = 0: Linear_projection + dropout = 0 (everything prior has a dropout = 0.2)
- Learned Linear Projection from the genePT embeddings to the scGPT embedding space prior to scGPT finetuning
- Linear Projection + Learned weights for each modality embedding (scGPT counts, scGPT tokens, genePT tokens)
- Separate transformer encoders for biology and language modalities

### A.3 Mapping of GO Term Annotations

Examples of post-processing the GO gene annotations.

**TABLE 22.** Concatenating GO Annotations. Examples for the FOSB gene.

| Annotation Type | Annotation Example | Concatenated String (*this gets embedded using an LLM*) |
| --- | --- | --- |
| **Gene Ontology** - Molecular Function | • DNA-binding transcription factor activity, RNA polymerase II-specific<br>• DNA-binding transcription activator activity, RNA polymerase II-specific<br>• DNA binding<br>• protein binding<br>• sequence-specific double-stranded DNA binding<br>• ... | **Molecular Function:** DNA-binding transcription factor activity, RNA polymerase II-specific, DNA-binding transcription activator activity, RNA polymerase II-specify, DNA binding, protein binding, sequence-specific double-stranded DNA binding, RNA polymerase II cis-regulatory region sequence-specific DNA binding |
| **Gene Ontology** - Cellular Component | • nucleus<br>• nucleoplasm<br>• cytosol<br>• intracellular membrane-bounded organelle<br>• chromatin | **Cellular Component:** nucleus, nucleoplasm, cytosol, intraceelular membrane-bounded organelle, chromatin |
| **Gene Ontology** - Biological Process | • negative regulation of transcription by RNA polymerase II<br>• response to amphetamine<br>• transcription by RNA polymerase II<br>• ... | **Biological Process:** negative regulation of transcription by RNA polymerase II, response to amphetamine, transcription by RNA polymerase II, female pregnancy .. |
| **Gene Ontology** - all | aggregation of all above | **Molecular Function:** DNA-binding transcription factor activity, RNA polymerase II-specific, DNA-binding transcription activator activity, ...;<br>**Cellular Component:** nucleus, nucleoplasm, cytosol, intraceelular membrane-bounded organelle, chromatin;<br>**Biological Process:** negative regulation of transcription by RNA polymerase II, response to amphetamine, transcription by RNA polymerase II, female pregnancy .. |

### A.4 UMAP Projections

**FIGURE 11.**
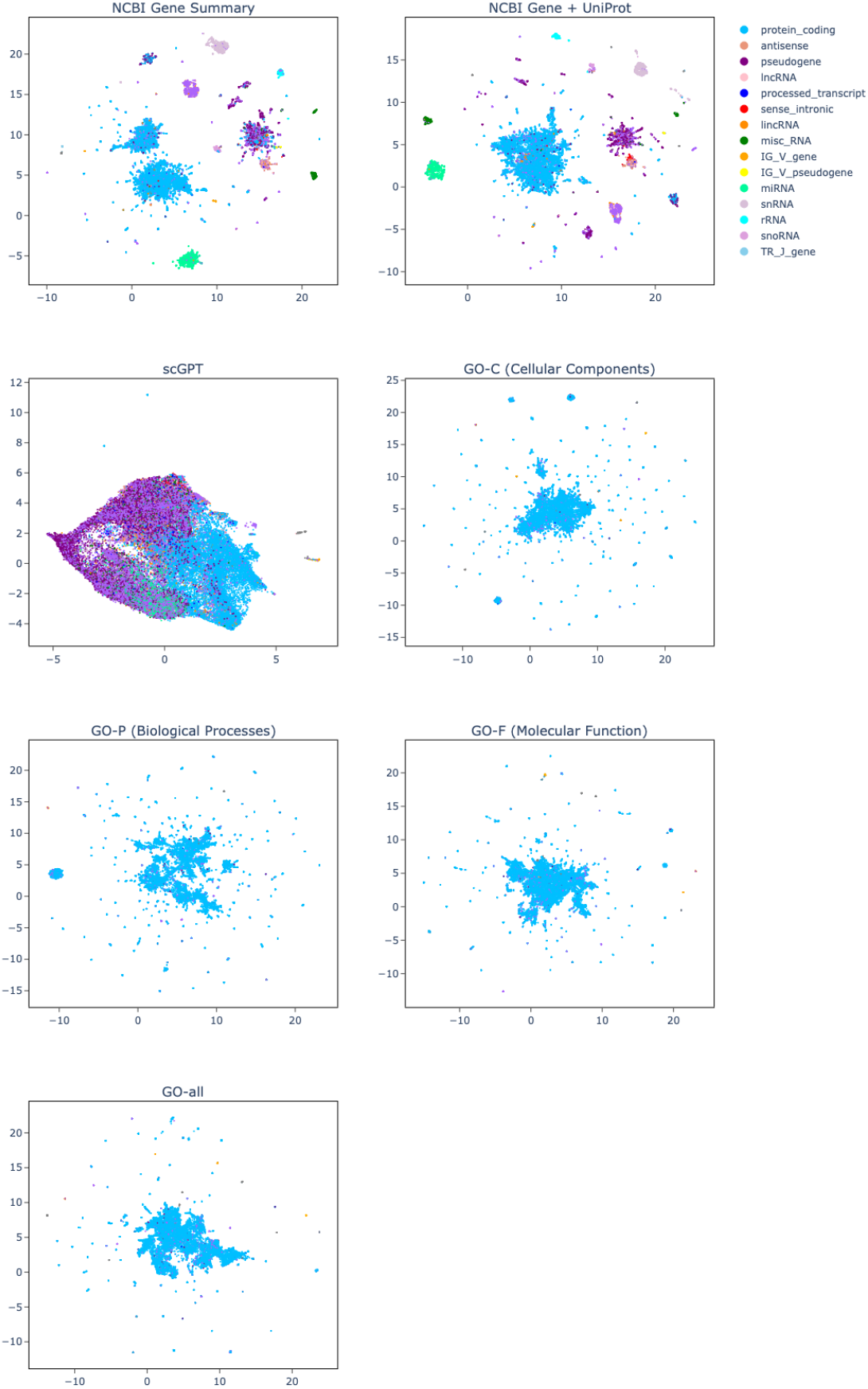
UMAP Projections of gene textual annotations embeddings, GPT-3.5, averaging GO Annotations. All annotations besides NCBI Gene + UniProt Summaries were embedded with *GPT-3*.*5-ada* embedding model. The NCBI Gene + UniProt Summaries was embedded with *GPT-3*.*5-text-embedding-3-large* model. The GO annotations used the **average** method. Each color corresponds to a different gene functionality.

**FIGURE 12.**
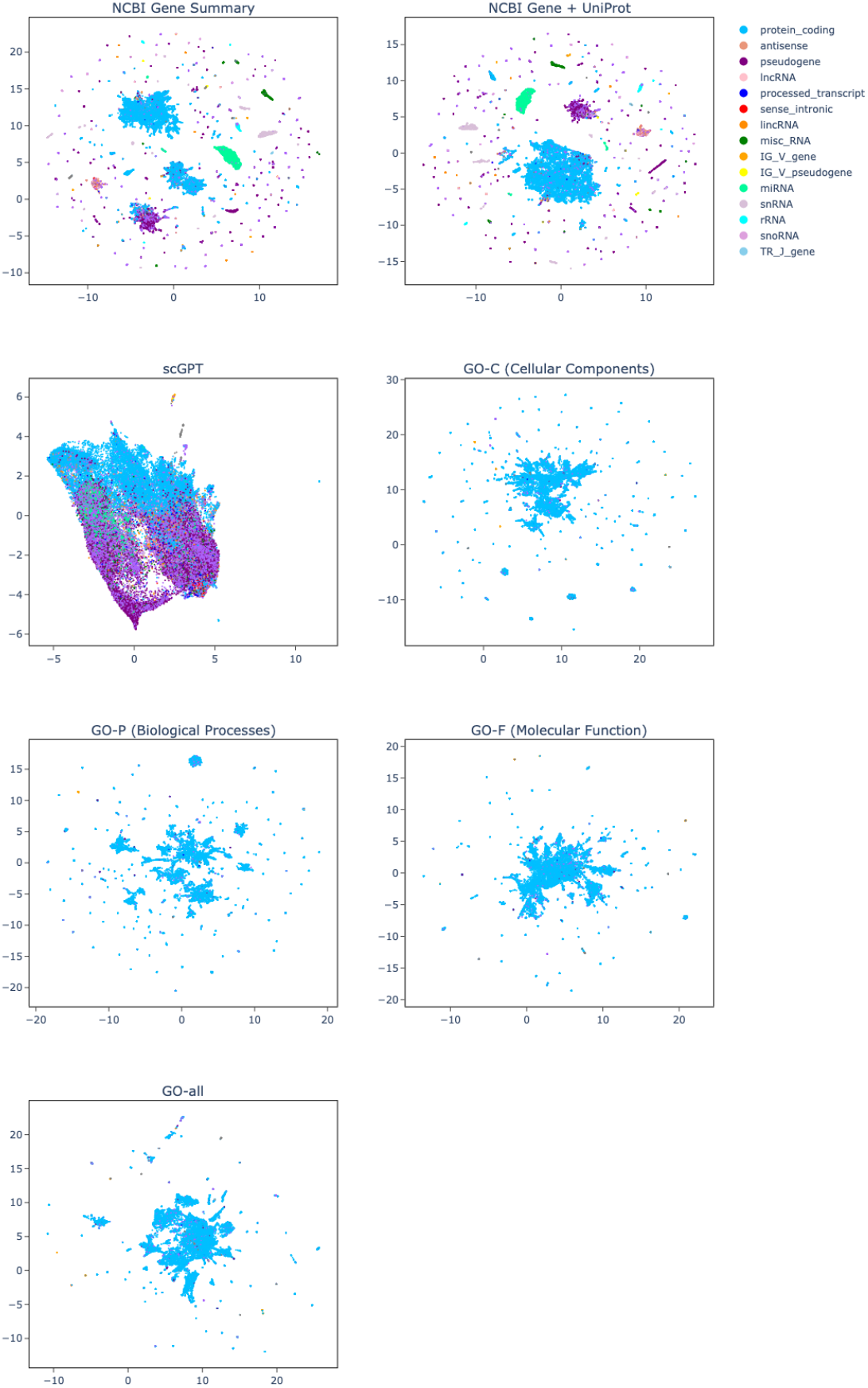
UMAP Projections of gene textual annotations embeddings, LLAMA-3.1-8b, concatenating GO Annotations All annotations were embedded using the *LLAMA-3*.*1-8b embedding model*. The GO annotations used the **concatenationn** method. Each color corresponds to a different gene functionality.

**FIGURE 13.**
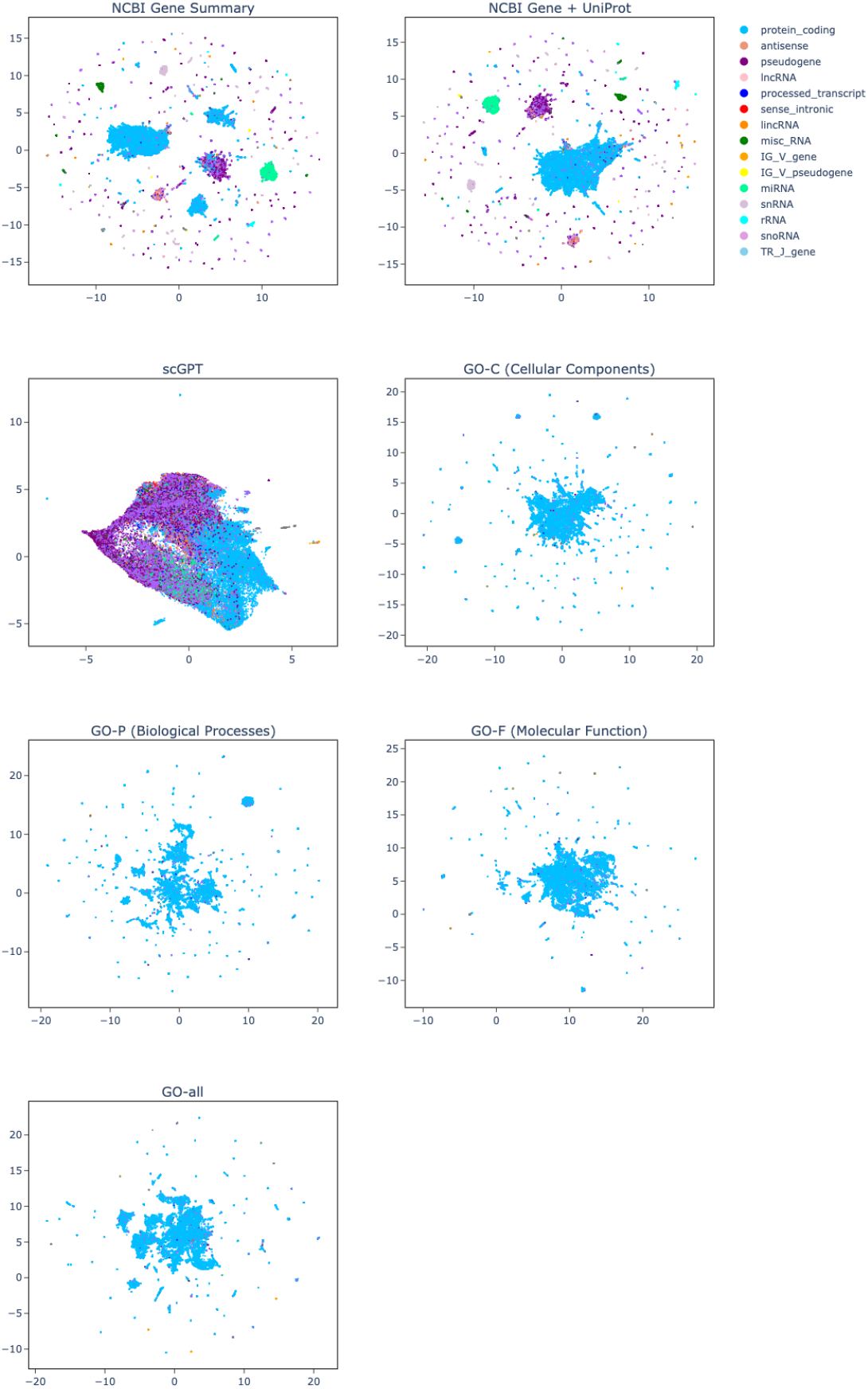
UMAP Projections of gene textual annotations embeddings, LLAMA-3.1-70b, concatenating GO Annotations. All annotations were embedded using the *LLAMA-3*.*1-70b embedding model*. The GO annotations used the **concatenationn** method. Each color corresponds to a different gene functionality.

### A.5 Effect of constructing GO Terms

#### A.5.1 Adamson Dataset

**TABLE 23.** GO terms avg vs concat, Adamson Dataset, Top20 Genes.

| Model | Pearson<br>Corr<br>($\Delta$)<sub>Top20</sub> ↑ | MSE<sub>Top20</sub> ↓ | Model | Pearson<br>Corr<br>($\Delta$)<sub>Top20</sub> ↑ | MSE<sub>Top20</sub> ↓ |
| --- | --- | --- | --- | --- | --- |
| scGPT | 0.782 ± 0.02 | 0.135 ± 0.01 | scGPT | 0.782 ± 0.02 | 0.135 ± 0.01 |
| scGenePT<sup>conc</sup><sub>GO-F</sub> | **0.785 ± 0.02** | **0.128 ± 0.00** | scGenePT<sup>avg</sup><sub>GO-F</sub> | 0.783 ± 0.01 | 0.130 ± 0.00 |
| scGenePT<sup>conc</sup><sub>GO-C</sub> | **0.789 ± 0.01** | **0.127 ± 0.00** | scGenePT<sup>avg</sup><sub>GO-C</sub> | 0.788 ± 0.02 | **0.127 ± 0.01** |
| scGenePT<sup>conc</sup><sub>GO-P</sub> | **0.791 ± 0.01** | **0.125 ± 0.00** | scGenePT<sup>avg</sup><sub>GO-P</sub> | 0.789 ± 0.02 | 0.126 ± 0.01 |
| scGenePT<sup>conc</sup><sub>GO-all</sub> | **0.787 ± 0.02** | **0.127 ± 0.01** | scGenePT<sup>avg</sup><sub>GO-all</sub> | 0.784 ± 0.02 | 0.129 ± 0.00 |

**TABLE 24.** GO terms avg vs concat, Adamson Dataset, All Genes.

| Model | Pearson<br>Corr<br>($\Delta$)<sub>All</sub> ↑ | MSE<sub>All</sub> ↓ | Model | Pearson<br>Corr<br>($\Delta$)<sub>All</sub> ↑ | MSE<sub>All</sub> ↓ |
| --- | --- | --- | --- | --- | --- |
| scGPT | 0.589 ± 0.03 | 0.00672 ± 0.00 | scGPT | 0.589 ± 0.03 | 0.00672 ± 0.00 |
| scGenePT<sup>conc</sup><sub>GO-F</sub> | **0.611 ± 0.03** | **0.00640 ± 0.00** | scGenePT<sup>avg</sup><sub>GO-F</sub> | 0.598 ± 0.04 | 0.00660 ± 0.00 |
| scGenePT<sup>conc</sup><sub>GO-C</sub> | 0.609 ± 0.03 | 0.00645 ± 0.00 | scGenePT<sup>avg</sup><sub>GO-C</sub> | **0.616 ± 0.03** | **0.00621 ± 0.00** |
| scGenePT<sup>conc</sup><sub>GO-P</sub> | **0.623 ± 0.02** | **0.00622 ± 0.00** | scGenePT<sup>avg</sup><sub>GO-P</sub> | 0.618 ± 0.03 | 0.00630 ± 0.00 |
| scGenePT<sup>conc</sup><sub>GO-all</sub> | 0.605 ± 0.03 | 0.00641 ± 0.00 | scGenePT<sup>avg</sup><sub>GO-all</sub> | **0.612 ± 0.03** | **0.00629 ± 0.00** |

#### A.5.2 Norman Dataset

**TABLE 25.** GO terms avg vs concat, Norman Dataset, Top20 Genes.

| Model | Pearson<br>Corr<br>($\Delta$)<sub>Top20</sub> ↑ | MSE<sub>Top20</sub> ↓ | Model | Pearson<br>Corr<br>($\Delta$)<sub>Top20</sub> ↑ | MSE<sub>Top20</sub> ↓ |
| --- | --- | --- | --- | --- | --- |
| scGPT | 0.534 ± 0.03 | 0.223 ± 0.02 | scGPT | 0.534 ± 0.03 | 0.223 ± 0.02 |
| scGenePT<sup>conc</sup><sub>GO-F</sub> | **0.554 ± 0.02** | **0.216 ± 0.02** | scGenePT<sup>avg</sup><sub>GO-F</sub> | 0.557 ± 0.03 | 0.217 ± 0.03 |
| scGenePT<sup>conc</sup><sub>GO-C</sub> | **0.543 ± 0.02** | 0.219 ± 0.01 | scGenePT<sup>avg</sup><sub>GO-C</sub> | 0.547 ± 0.02 | **0.216 ± 0.03** |
| scGenePT<sup>conc</sup><sub>GO-P</sub> | 0.550 ± 0.02 | **0.220 ± 0.02** | scGenePT<sup>avg</sup><sub>GO-P</sub> | **0.553 ± 0.03** | 0.223 ± 0.02 |
| scGenePT<sup>conc</sup><sub>GO-all</sub> | **0.554 ± 0.02** | **0.209 ± 0.02** | scGenePT<sup>avg</sup><sub>GO-all</sub> | 0.689 ± 0.02 | 0.216 ± 0.02 |

**TABLE 26.** GO terms avg vs concat, Norman Dataset, All Genes.

| Model | Pearson Corr<br>($\Delta$)<sub>All</sub> ↑ | MSE<sub>All</sub> ↓ | Model | Pearson Corr<br>($\Delta$)<sub>All</sub> ↑ | MSE<sub>All</sub> ↓ |
| --- | --- | --- | --- | --- | --- |
| scGPT | $0.534 \pm 0.03$ | $0.00421 \pm 0.00$ | scGPT | $0.534 \pm 0.03$ | $0.00421 \pm 0.00$ |
| scGenePT<sup>conc</sup><sub>GO-F</sub> | $0.554 \pm 0.02$ | **$0.00405 \pm 0.00$** | scGenePT<sup>avg</sup><sub>GO-F</sub> | **$0.557 \pm 0.03$** | $0.00407 \pm 0.00$ |
| scGenePT<sup>conc</sup><sub>GO-C</sub> | $0.543 \pm 0.02$ | **$0.00412 \pm 0.00$** | scGenePT<sup>avg</sup><sub>GO-C</sub> | **$0.547 \pm 0.02$** | $0.00418 \pm 0.00$ |
| scGenePT<sup>conc</sup><sub>GO-P</sub> | $0.550 \pm 0.02$ | **$0.00405 \pm 0.00$** | scGenePT<sup>avg</sup><sub>GO-P</sub> | **$0.553 \pm 0.03$** | $0.00409 \pm 0.00$ |
| scGenePT<sup>conc</sup><sub>GO-all</sub> | **$0.554 \pm 0.02$** | **$0.00407 \pm 0.00$** | scGenePT<sup>avg</sup><sub>GO-all</sub> | $0.549 \pm 0.02$ | $0.00410 \pm 0.00$ |

### A.6 genePT vs scGenePT, Adamson Dataset

**FIGURE 14.**
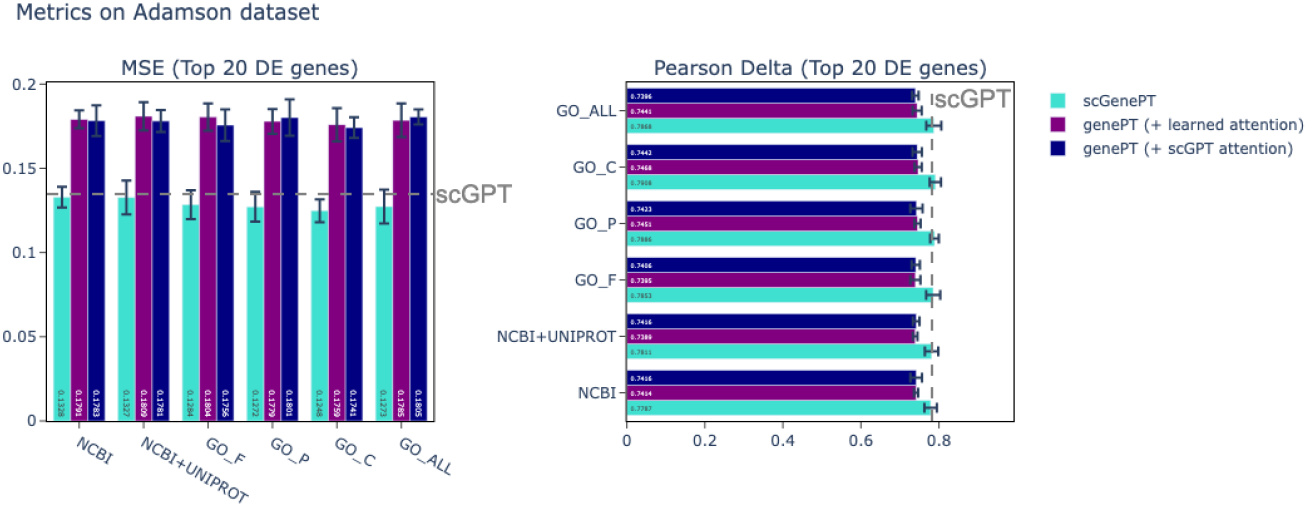
Comparison of performance of scGenePT with different language embeddings and language embeddings alone on the test split of the Adamson dataset.

**TABLE 27.** genePT vs scGenePT, Adamson Dataset, **Pearson Corr Δ**_**Top20**_.

| Model | Pearson Corr<br>$\Delta_{\text{Top20}}$ ↑ | Model | Pearson Corr<br>$\Delta_{\text{Top20}}$ ↑<br>(learned attention) | Pearson Corr<br>$\Delta_{\text{Top20}}$ ↑<br>(scGPT attention) |
| --- | --- | --- | --- | --- |
| scGPT | $0.782 \pm 0.02$ | scGPT | $0.782 \pm 0.02$ | $0.782 \pm 0.02$ |
| scGenePT<sub>NCBI</sub> | **$0.779 \pm 0.02$** | genePT<sub>NCBI</sub> | $0.741 \pm 0.00$ | $0.742 \pm 0.02^*$ |
| scGenePT<sub>NCBI+UniProt</sub> | **$0.781 \pm 0.02$** | genePT<sub>NCBI+UniProt</sub> | $0.739 \pm 0.00$ | $0.742 \pm 0.00^*$ |
| scGenePT<sub>GO-F</sub> | **$0.785 \pm 0.01$** | genePT<sub>GO-F</sub> | $0.740 \pm 0.00$ | $0.741 \pm 0.01^*$ |
| scGenePT<sub>GO-C</sub> | **$0.791 \pm 0.01$** | genePT<sub>GO-C</sub> | $0.747 \pm 0.00^*$ | $0.744 \pm 0.01$ |
| scGenePT<sub>GO-P</sub> | **$0.789 \pm 0.01$** | genePT<sub>GO-P</sub> | $0.745 \pm 0.00^*$ | $0.742 \pm 0.02$ |
| scGenePT<sub>GO-all</sub> | **$0.787 \pm 0.02$** | genePT<sub>GO-all</sub> | $0.744 \pm 0.01^*$ | $0.740 \pm 0.00$ |

**TABLE 28.** genePT vs scGenePT, Adamson Dataset, **Pearson Corr Δ**_**All**_.

| Model | Pearson Corr ($\Delta_{All}$) ↑ | Model | Pearson Corr ($\Delta_{All}$) ↑ (learned attention) | Pearson Corr ($\Delta_{All}$) ↑ (scGPT attention) |
| --- | --- | --- | --- | --- |
| scGPT | $0.589 \pm 0.03$ | scGPT | $0.589 \pm 0.03$ | $0.589 \pm 0.03$ |
| scGenePT<sub>NCBI</sub> | **$0.606 \pm 0.03$** | genePT<sub>NCBI</sub> | $0.344 \pm 0.02^*$ | $0.340 \pm 0.03$ |
| scGenePT<sub>NCBI+UniProt</sub> | **$0.617 \pm 0.03$** | genePT<sub>NCBI+UniProt</sub> | $0.331 \pm 0.01$ | $0.339 \pm 0.02^*$ |
| scGenePT<sub>GO-F</sub> | **$0.611 \pm 0.03$** | genePT<sub>GO-F</sub> | $0.334 \pm 0.02$ | $0.339 \pm 0.01^*$ |
| scGenePT<sub>GO-C</sub> | **$0.623 \pm 0.03$** | genePT<sub>GO-C</sub> | $0.350 \pm 0.00^*$ | $0.342 \pm 0.03$ |
| scGenePT<sub>GO-P</sub> | **$0.609 \pm 0.03$** | genePT<sub>GO-P</sub> | $0.350 \pm 0.01^*$ | $0.337 \pm 0.03$ |
| scGenePT<sub>GO-all</sub> | **$0.605 \pm 0.03$** | genePT<sub>GO-all</sub> | $0.351 \pm 0.03^*$ | $0.337 \pm 0.01$ |

**TABLE 29.** genePT vs scGenePT, Adamson Dataset. **MSE**_**Top20**_.

| Model | MSE<sub>Top20</sub> ↓ | Model | MSE<sub>Top20</sub> ↓ (learned attention) | MSE<sub>Top20</sub> ↓ (scGPT attention) |
| --- | --- | --- | --- | --- |
| scGPT | $0.135 \pm 0.01$ | scGPT | $0.135 \pm 0.01$ | $0.135 \pm 0.01$ |
| scGenePT<sub>NCBI</sub> | **$0.133 \pm 0.00$** | genePT<sub>NCBI</sub> | $0.179 \pm 0.00$ | $0.178 \pm 0.00^*$ |
| scGenePT<sub>NCBI+UniProt</sub> | **$0.133 \pm 0.01$** | genePT<sub>NCBI+UniProt</sub> | $0.181 \pm 0.00$ | $0.178 \pm 0.00^*$ |
| scGenePT<sub>GO-F</sub> | **$0.128 \pm 0.00$** | genePT<sub>GO-F</sub> | $0.180 \pm 0.00$ | $0.176 \pm 0.00^*$ |
| scGenePT<sub>GO-C</sub> | **$0.125 \pm 0.00$** | genePT<sub>GO-C</sub> | $0.176 \pm 0.01$ | $0.174 \pm 0.00^*$ |
| scGenePT<sub>GO-P</sub> | **$0.127 \pm 0.00$** | genePT<sub>GO-P</sub> | $0.178 \pm 0.00^*$ | $0.180 \pm 0.01$ |
| scGenePT<sub>GO-all</sub> | **$0.127 \pm 0.01$** | genePT<sub>GO-all</sub> | $0.179 \pm 0.01^*$ | $0.181 \pm 0.00$ |

**TABLE 30.** genePT vs scGenePT, Adamson Dataset. **MSE**_**All**_.

| Model | MSE<sub>All</sub> ↓ | Model | MSE<sub>All</sub> ↓ (learned attention) | MSE<sub>All</sub> ↓ (scGPT attention) |
| --- | --- | --- | --- | --- |
| scGPT | $0.00672 \pm 0.01$ | scGPT | $0.135 \pm 0.01$ | $0.135 \pm 0.01$ |
| scGenePT<sub>NCBI</sub> | **$0.00654 \pm 0.00$** | genePT<sub>NCBI</sub> | $0.02461 \pm 0.00^*$ | $0.02462 \pm 0.00$ |
| scGenePT<sub>NCBI+UniProt</sub> | **$0.00648 \pm 0.00$** | genePT<sub>NCBI+UniProt</sub> | $0.02436 \pm 0.00^*$ | $0.02437 \pm 0.00$ |
| scGenePT<sub>GO-F</sub> | **$0.00640 \pm 0.00$** | genePT<sub>GO-F</sub> | $0.02452 \pm 0.00$ | $0.02438 \pm 0.00^*$ |
| scGenePT<sub>GO-C</sub> | **$0.00622 \pm 0.00$** | genePT<sub>GO-C</sub> | $0.02454 \pm 0.00$ | $0.02451 \pm 0.00^*$ |
| scGenePT<sub>GO-P</sub> | **$0.00645 \pm 0.00$** | genePT<sub>GO-P</sub> | $0.02466 \pm 0.00$ | $0.02456 \pm 0.00^*$ |
| scGenePT<sub>GO-all</sub> | **$0.00641 \pm 0.00$** | genePT<sub>GO-all</sub> | $0.02459 \pm 0.00$ | $0.02450 \pm 0.00^*$ |

### A.7 Training Parameters

**TABLE 31.**
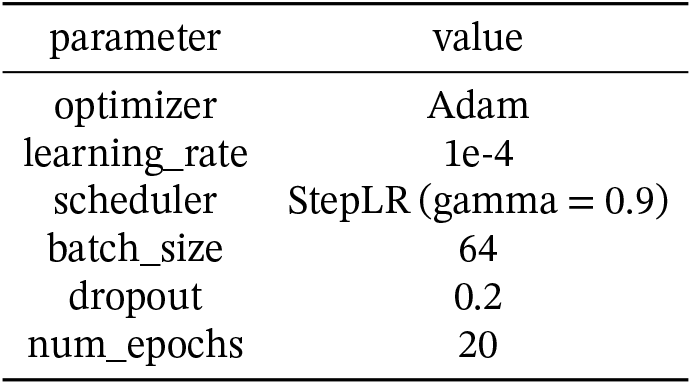
Training Parameters.

### A.8 Effect of the language embedding choice

#### A.8.1 Norman Dataset

**FIGURE 15.**
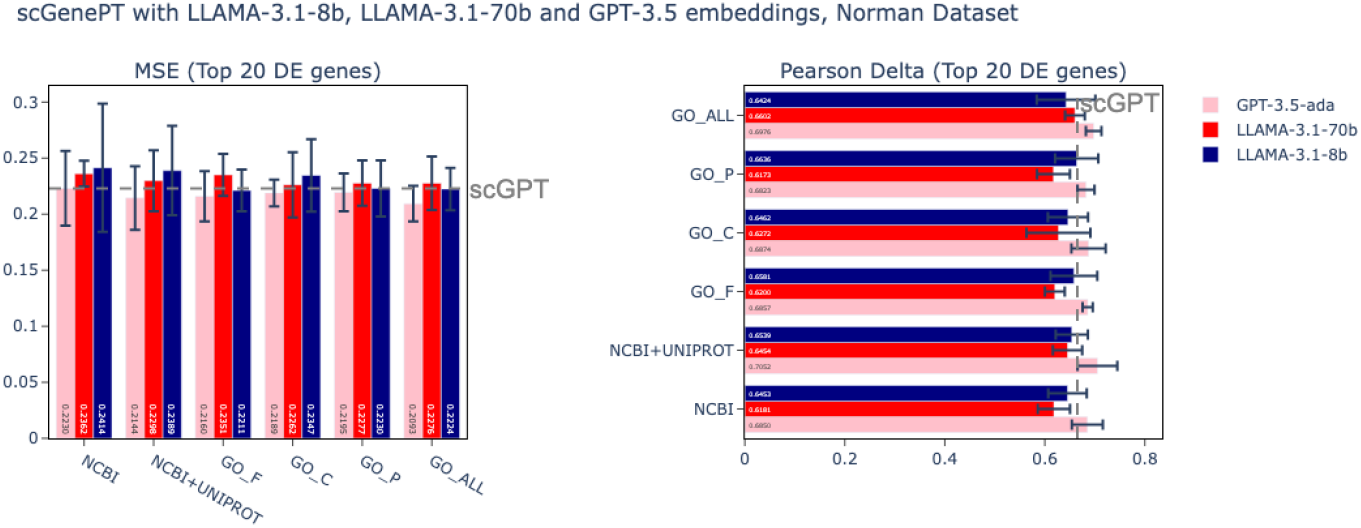
**Pearson Correlation Δ**_**Top20**_ for scGenePT using GPT-3.5, LLAMA-3.1-8b, and LLAMA-3.1-70b embedding models, Norman Dataset

**TABLE 32.** **Pearson Correlation Δ**_**Top20**_ ↑ Norman Dataset, Different Embedding Models.

| Model | GPT-3.5 | LLAMA-3.1-70b | LLAMA-3.1-8b |
| --- | --- | --- | --- |
| scGPT | $0.665 \pm 0.01$ | $0.665 \pm 0.01$ | $0.665 \pm 0.01$ |
| scGenePT<sub>NCBI</sub> | $0.685 \pm 0.03$ | $0.618 \pm 0.03$ | $0.645 \pm 0.04$ |
| scGenePT<sub>NCBI+UniProt</sub> | $0.705 \pm 0.04$ | $0.645 \pm 0.03$ | $0.654 \pm 0.04$ |
| scGenePT<sub>GO-F</sub> | $0.686 \pm 0.01$ | $0.620 \pm 0.06$ | $0.658 \pm 0.05$ |
| scGenePT<sub>GO-C</sub> | $0.687 \pm 0.03$ | $0.627 \pm 0.03$ | $0.646 \pm 0.04$ |
| scGenePT<sub>GO-P</sub> | $0.682 \pm 0.02$ | $0.617 \pm 0.03$ | $0.664 \pm 0.04$ |
| scGenePT<sub>GO-all</sub> | $0.698 \pm 0.02$ | $0.660 \pm 0.02$ | $0.642 \pm 0.06$ |

**TABLE 33.**
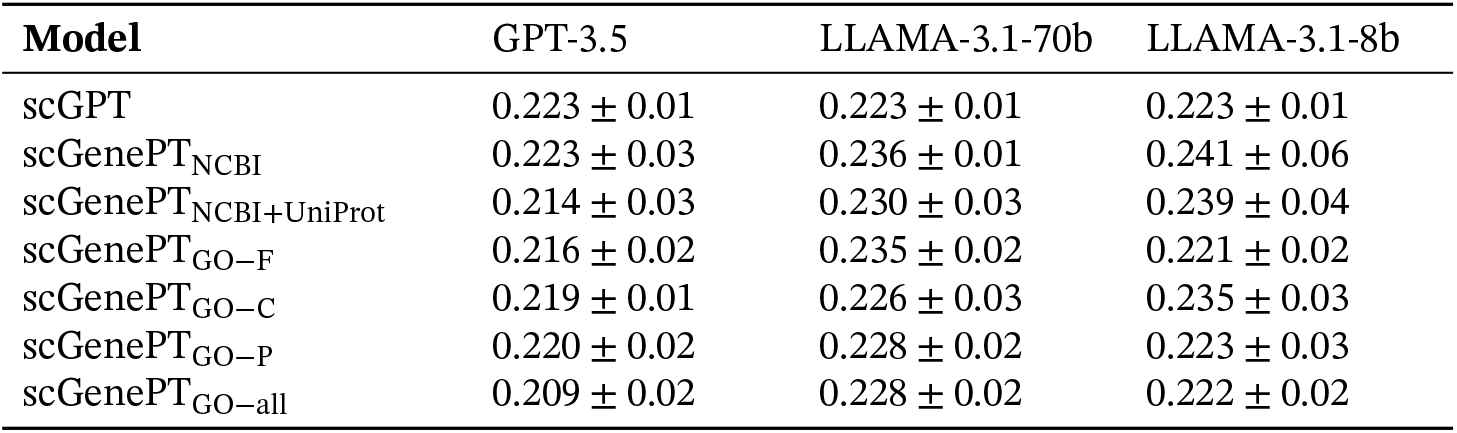
**MSE Δ**_**Top20**_ ↓ Norman Dataset, Different Embedding Models.

**TABLE 34.**
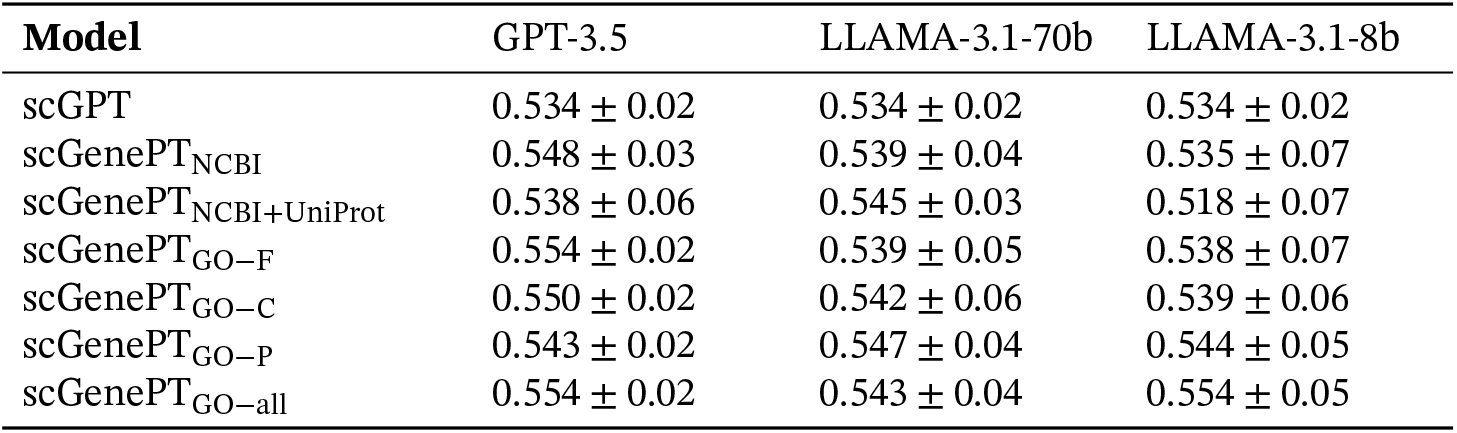
**Pearson Correlation Δ**_**All**_↑ Norman Dataset, Different Embedding Models.

**TABLE 35.** **MSE** _**All**_ ↓ Norman Dataset, Different Embedding Models.

| Model | GPT-3.5 | LLAMA-3.1-70b | LLAMA-3.1-8b |
| --- | --- | --- | --- |
| scGPT | $0.00421 \pm 0.00$ | $0.00421 \pm 0.00$ | $0.0421 \pm 0.00$ |
| scGenePT<sub>NCBI</sub> | $0.00415 \pm 0.00$ | $0.00416 \pm 0.03$ | $0.00423 \pm 0.00$ |
| scGenePT<sub>NCBI+UniProt</sub> | $0.00413 \pm 0.00$ | $0.00413 \pm 0.00$ | $0.00427 \pm 0.00$ |
| scGenePT<sub>GO-F</sub> | $0.00405 \pm 0.00$ | $0.00410 \pm 0.00$ | $0.00404 \pm 0.00$ |
| scGenePT<sub>GO-C</sub> | $0.00405 \pm 0.00$ | $0.00406 \pm 0.00$ | $0.00413 \pm 0.00$ |
| scGenePT<sub>GO-P</sub> | $0.00412 \pm 0.00$ | $0.00406 \pm 0.00$ | $0.00412 \pm 0.00$ |
| scGenePT<sub>GO-all</sub> | $0.00407 \pm 0.00$ | $0.00416 \pm 0.00$ | $0.00406 \pm 0.00$ |

#### A.8.2 Adamson Dataset

**FIGURE 16.**
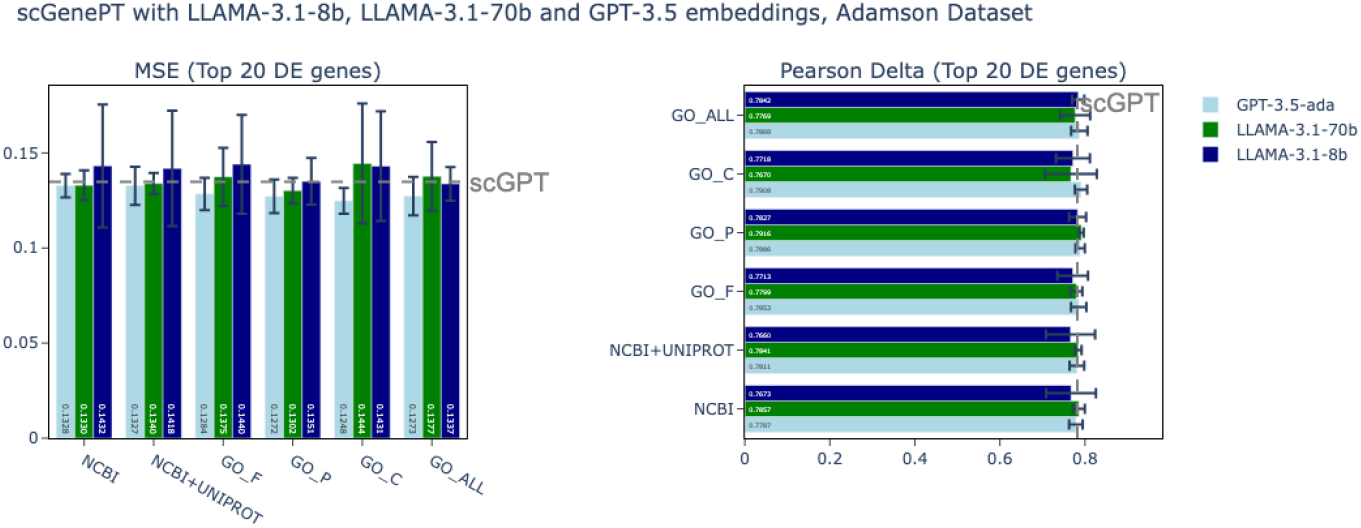
**Pearson Correlation Δ**_**Top20**_ for scGenePT using GPT-3.5, LLAMA-3.1-8b, and LLAMA-3.1-70b embedding models, Adamson Dataset

**TABLE 36.** **Pearson CorrelationΔ**_**Top20**_ ↑Adamson Dataset, Different Embedding Models.

| Model | GPT-3.5 | LLAMA-3.1-70b | LLAMA-3.1-8b |
| --- | --- | --- | --- |
| scGPT | $0.782 \pm 0.02$ | $0.782 \pm 0.02$ | $0.782 \pm 0.02$ |
| scGenePT<sub>NCBI</sub> | $0.779 \pm 0.02$ | $0.786 \pm 0.01$ | $0.767 \pm 0.06$ |
| scGenePT<sub>NCBI+UniProt</sub> | $0.781 \pm 0.02$ | $0.784 \pm 0.00$ | $0.766 \pm 0.06$ |
| scGenePT<sub>GO-F</sub> | $0.785 \pm 0.02$ | $0.780 \pm 0.00$ | $0.771 \pm 0.04$ |
| scGenePT<sub>GO-C</sub> | $0.789 \pm 0.01$ | $0.792 \pm 0.00$ | $0.783 \pm 0.02$ |
| scGenePT<sub>GO-P</sub> | $0.791 \pm 0.01$ | $0.767 \pm 0.06$ | $0.772 \pm 0.04$ |
| scGenePT<sub>GO-all</sub> | $0.787 \pm 0.02$ | $0.777 \pm 0.04$ | $0.784 \pm 0.01$ |

**TABLE 37.** **MSE** _**Top20**_ ↓ Adamson Dataset, Different Embedding Models.

| Model | GPT-3.5 | LLAMA-3.1-70b | LLAMA-3.1-8b |
| --- | --- | --- | --- |
| scGPT | $0.135 \pm 0.01$ | $0.135 \pm 0.01$ | $0.135 \pm 0.01$ |
| scGenePT<sub>NCBI</sub> | $0.133 \pm 0.00$ | $0.133 \pm 0.00$ | $0.143 \pm 0.03$ |
| scGenePT<sub>NCBI+UniProt</sub> | $0.133 \pm 0.01$ | $0.134 \pm 0.00$ | $0.142 \pm 0.03$ |
| scGenePT<sub>GO-F</sub> | $0.128 \pm 0.02$ | $0.137 \pm 0.02$ | $0.144 \pm 0.03$ |
| scGenePT<sub>GO-C</sub> | $0.127 \pm 0.00$ | $0.130 \pm 0.00$ | $0.135 \pm 0.01$ |
| scGenePT<sub>GO-P</sub> | $0.125 \pm 0.00$ | $0.144 \pm 0.03$ | $0.143 \pm 0.03$ |
| scGenePT<sub>GO-all</sub> | $0.127 \pm 0.01$ | $0.138 \pm 0.02$ | $0.134 \pm 0.00$ |

**TABLE 38.** **Pearson Correlation Δ**_**All**_ ↑Adamson Dataset, Different Embedding Models.

| **Model** | GPT-3.5 | LLAMA-3.1-70b | LLAMA-3.1-8b |
| --- | --- | --- | --- |
| scGPT | $0.589 \pm 0.03$ | $0.589 \pm 0.03$ | $0.589 \pm 0.03$ |
| scGenePT<sub>NCBI</sub> | $0.606 \pm 0.03$ | $0.603 \pm 0.04$ | $0.597 \pm 0.05$ |
| scGenePT<sub>NCBI+UniProt</sub> | $0.607 \pm 0.03$ | $0.615 \pm 0.02$ | $0.599 \pm 0.05$ |
| scGenePT<sub>GO-F</sub> | $0.611 \pm 0.03$ | $0.605 \pm 0.04$ | $0.603 \pm 0.04$ |
| scGenePT<sub>GO-C</sub> | $0.609 \pm 0.03$ | $0.615 \pm 0.02$ | $0.607 \pm 0.04$ |
| scGenePT<sub>GO-P</sub> | $0.623 \pm 0.02$ | $0.592 \pm 0.05$ | $0.596 \pm 0.04$ |
| scGenePT<sub>GO-all</sub> | $0.605 \pm 0.03$ | $0.598 \pm 0.03$ | $0.609 \pm 0.03$ |

**TABLE 39.**
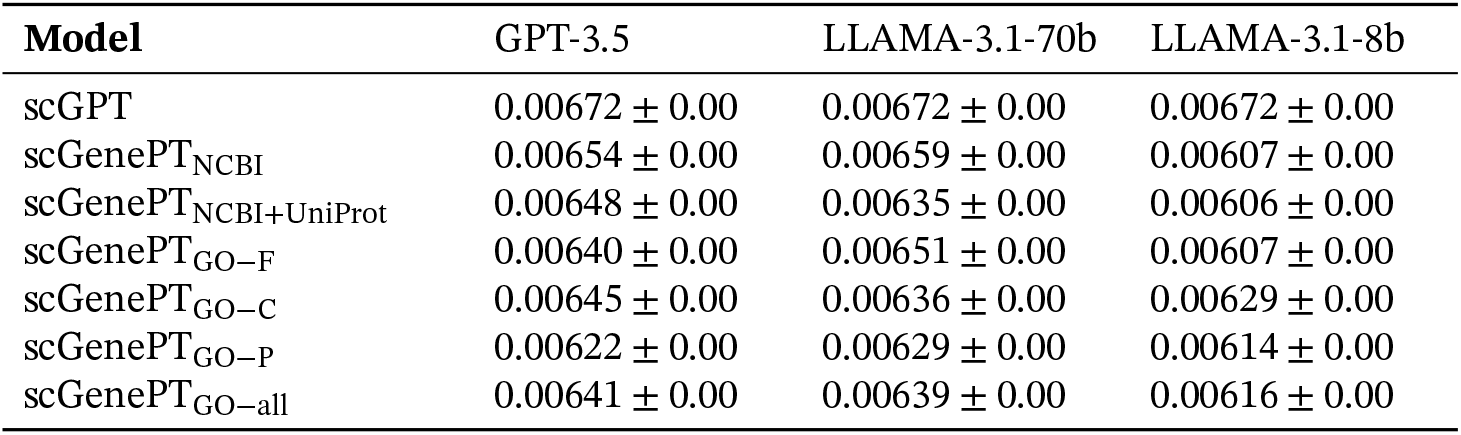
**MSE** _**All**_ ↓ Norman Dataset, Different Embedding Models.

| **Model** | GPT-3.5 | LLAMA-3.1-70b | LLAMA-3.1-8b |
| --- | --- | --- | --- |
| scGPT | $0.00672 \pm 0.00$ | $0.00672 \pm 0.00$ | $0.00672 \pm 0.00$ |
| scGenePT<sub>NCBI</sub> | $0.00654 \pm 0.00$ | $0.00659 \pm 0.00$ | $0.00607 \pm 0.00$ |
| scGenePT<sub>NCBI+UniProt</sub> | $0.00648 \pm 0.00$ | $0.00635 \pm 0.00$ | $0.00606 \pm 0.00$ |
| scGenePT<sub>GO-F</sub> | $0.00640 \pm 0.00$ | $0.00651 \pm 0.00$ | $0.00607 \pm 0.00$ |
| scGenePT<sub>GO-C</sub> | $0.00645 \pm 0.00$ | $0.00636 \pm 0.00$ | $0.00629 \pm 0.00$ |
| scGenePT<sub>GO-P</sub> | $0.00622 \pm 0.00$ | $0.00629 \pm 0.00$ | $0.00614 \pm 0.00$ |
| scGenePT<sub>GO-all</sub> | $0.00641 \pm 0.00$ | $0.00639 \pm 0.00$ | $0.00616 \pm 0.00$ |

## References

[1] A Vaswani. Attention is all you need. Advances in Neural Information Processing Systems, 2017.

[2] Yinhan Liu. Roberta: A robustly optimized bert pretraining approach. arXiv preprint arXiv:1907.11692, 2019.

[3] Alec Radford, Jeffrey Wu, Rewon Child, David Luan, Dario Amodei, Ilya Sutskever, et al. Language models are unsupervised multitask learners. OpenAI blog, 1(8):9, 2019.

[4] Tom B Brown. Language models are few-shot learners. arXiv preprint arXiv:2005.14165, 2020.

[5] Hugo Touvron, Thibaut Lavril, Gautier Izacard, Xavier Martinet, Marie-Anne Lachaux, Timothée Lacroix, Baptiste Rozière, Naman Goyal, Eric Hambro, Faisal Azhar, et al. Llama: Open and efficient foundation language models. arXiv preprint arXiv:2302.13971, 2023.

[6] Abhimanyu Dubey, Abhinav Jauhri, Abhinav Pandey, Abhishek Kadian, Ahmad Al-Dahle, Aiesha Letman, Akhil Mathur, Alan Schelten, Amy Yang, Angela Fan, et al. The llama 3 herd of models. arXiv preprint arXiv:2407.21783, 2024.

[7] Alexey Dosovitskiy. An image is worth 16×16 words: Transformers for image recognition at scale. arXiv preprint arXiv:2010.11929, 2020.

[8] Haotian Cui, Chloe Wang, Hassaan Maan, Kuan Pang, Fengning Luo, Nan Duan, and Bo Wang. scgpt: toward building a foundation model for single-cell multi-omics using generative ai. Nature Methods, pages 1–11, 2024.

[9] Yiqun Chen and James Zou. Genept: A simple but effective foundation model for genes and cells built from chatgpt. bioRxiv, 2023.

[10] Yanay Rosen, Yusuf Roohani, Ayush Agrawal, Leon Samotorčan, Tabula Sapiens Consortium, Stephen R Quake, and Jure Leskovec. Universal cell embeddings: A foundation model for cell biology. bioRxiv, pages 2023–11, 2023.

[11] Christina V Theodoris, Ling Xiao, Anant Chopra, Mark D Chaffin, Zeina R Al Sayed, Matthew C Hill, Helene Mantineo, Elizabeth M Brydon, Zexian Zeng, X Shirley Liu, et al. Transfer learning enables predictions in network biology. Nature, 618(7965):616–624, 2023.

[12] CZI Single-Cell Biology, Shibla Abdulla, Brian Aevermann, Pedro Assis, Seve Bada-joz, Sidney M Bell, Emanuele Bezzi, Batuhan Cakir, Jim Chaffer, Signe Chambers, et al. Cz cellxgene discover: A single-cell data platform for scalable exploration, analysis and modeling of aggregated data. BioRxiv, pages 2023–10, 2023.

[13] Haiyang Bian, Yixin Chen, Xiaomin Dong, Chen Li, Minsheng Hao, Sijie Chen, Jinyi Hu, Maosong Sun, Lei Wei, and Xuegong Zhang. scmulan: a multitask generative pre-trained language model for single-cell analysis. In International Conference on Research in Computational Molecular Biology, pages 479–482. Springer, 2024.

[14] Yusuf Roohani, Kexin Huang, and Jure Leskovec. Predicting transcriptional outcomes of novel multigene perturbations with gears. Nature Biotechnology, 2023.

[15] Garth R Brown, Vichet Hem, Kenneth S Katz, Michael Ovetsky, Craig Wallin, Olga Ermolaeva, Igor Tolstoy, Tatiana Tatusova, Kim D Pruitt, Donna R Maglott, et al. Gene: a gene-centered information resource at ncbi. Nucleic acids research, 43 (D1):D36–D42, 2015.

[16] Uniprot: the universal protein knowledgebase in 2023. Nucleic acids research, 51 (D1):D523–D531, 2023.

[17] Michael Ashburner, Catherine A Ball, Judith A Blake, David Botstein, Heather Butler, J Michael Cherry, Allan P Davis, Kara Dolinski, Selina S Dwight, Janan T Eppig, et al. Gene ontology: tool for the unification of biology. Nature genetics, 25 (1):25–29, 2000.

[18] Suzi A Aleksander, James Balhoff, Seth Carbon, J Michael Cherry, Harold J Drabkin, Dustin Ebert, Marc Feuermann, Pascale Gaudet, Nomi L Harris, et al. The gene ontology knowledgebase in 2023. Genetics, 224(1):iyad031, 2023.

[19] Josh Achiam, Steven Adler, Sandhini Agarwal, Lama Ahmad, Ilge Akkaya, Floren-cia Leoni Aleman, Diogo Almeida, Janko Altenschmidt, Sam Altman, Shyamal Anadkat, et al. Gpt-4 technical report. arXiv preprint arXiv:2303.08774, 2023.

[20] Thomas M Norman, Max A Horlbeck, Joseph M Replogle, Alex Y Ge, Albert Xu, Marco Jost, Luke A Gilbert, and Jonathan S Weissman. Exploring genetic interaction manifolds constructed from rich single-cell phenotypes. Science, 365 (6455):786–793, 2019.

[21] Britt Adamson, Thomas M Norman, Marco Jost, Min Y Cho, James K Nuñez, Yuwen Chen, Jacqueline E Villalta, Luke A Gilbert, Max A Horlbeck, Marco Y Hein, et al. A multiplexed single-cell crispr screening platform enables systematic dissection of the unfolded protein response. Cell, 167(7):1867–1882, 2016.

[22] Michael Bereket and Theofanis Karaletsos. Modelling cellular perturbations with the sparse additive mechanism shift variational autoencoder. Advances in Neural Information Processing Systems, 36, 2024.

[23] Kaspar Märtens, Rory Donovan-Maiye, and Jesper Ferkinghoff-Borg. Enhancing generative perturbation models with llm-informed gene embeddings. In ICLR 2024 Workshop on Machine Learning for Genomics Explorations.

[24] Joseph M Replogle, Reuben A Saunders, Angela N Pogson, Jeffrey A Hussmann, Alexander Lenail, Alina Guna, Lauren Mascibroda, Eric J Wagner, Karen Adelman, Gila Lithwick-Yanai, et al. Mapping information-rich genotype-phenotype landscapes with genome-scale perturb-seq. Cell, 185(14):2559–2575, 2022.

[25] Moritz Schaefer, Peter Peneder, Daniel Malzl, Anna Hakobyan, Varun S Sharma, Thomas Krausgruber, Jörg Menche, Eleni Tomazou, and Christoph Bock. Joint embedding of transcriptomes and text enables interactive single-cell rna-seq data exploration via natural language. In ICLR 2024 Workshop on Machine Learning for Genomics Explorations.

[26] Jinhyuk Lee, Wonjin Yoon, Sungdong Kim, Donghyeon Kim, Sunkyu Kim, Chan Ho So, and Jaewoo Kang. Biobert: a pre-trained biomedical language representation model for biomedical text mining. Bioinformatics, 36(4):1234–1240, 2020.

[27] Guillaume Richard, Bernardo P de Almeida, Hugo Dalla-Torre, Christopher Blum, Lorenz Hexemer, Priyanka Pandey, Stefan Laurent, Marie P Lopez, Alexander Laterre, Maren Lang, et al. Chatnt: A multimodal conversational agent for dna, rna and protein tasks. bioRxiv, pages 2024–04, 2024.

[28] Daniel Levine, Sacha Lévy, Syed Asad Rizvi, Nazreen Pallikkavaliyaveetil, Xingyu Chen, David Zhang, Sina Ghadermarzi, Ruiming Wu, Zihe Zheng, Ivan Vrkic, et al. Cell2sentence: Teaching large language models the language of biology. bioRxiv, pages 2023–09, 2023.

[29] Hongyoon Choi, Jeongbin Park, Sumin Kim, Jiwon Kim, Dongjoo Lee, Sungwoo Bae, Haenara Shin, and Daeseung Lee. Cellama: Foundation model for single cell and spatial transcriptomics by cell embedding leveraging language model abilities. bioRxiv, pages 2024–05, 2024.

[30] Mohammad Lotfollahi, Anna Klimovskaia Susmelj, Carlo De Donno, Leon Hetzel, Yuge Ji, Ignacio L Ibarra, Sanjay R Srivatsan, Mohsen Naghipourfar, Riza M Daza, Beth Martin, et al. Predicting cellular responses to complex perturbations in high-throughput screens. Molecular systems biology, 19(6):e11517, 2023.

